# Joint loading in the presence of torsional deformities is overestimated unless gait adaptations are considered: a predictive simulation approach

**DOI:** 10.64898/2026.09.08.750035

**Authors:** Nicos Haralabidis, Elyse Passmore, Christopher Carty, Enrico De Pieri, Erich Rutz, Luca Modenese

**Author notes:** Corresponding author: Luca Modenese, School of Biomedical Engineering, University of New South Wales, Sydney, Australia, 2052.

## Abstract

Lower-limb torsional deformities have been shown to alter joint loading, although the findings vary between studies perhaps due to the simulation approaches applied. This study used predictive simulations to investigate how femoral neck anteversion (FNA) and external tibial torsion (ETT) influence hip and knee joint loading. Musculoskeletal models with altered FNA (1°–48°) and ETT (12°–53°), in isolation and combination, were created from a scaled adult model, and predictive walking (speed: 1.33 m/s) simulations were generated. Predictive simulations reproduced adaptations in hip rotation and foot-progression angle reported in individuals with torsional deformities. Hip and knee compressive, shear, and resultant contact forces were estimated, and multiple linear regressions quantified the independent associations of FNA and ETT with each outcome. Regressions accounted for 23%–86% of the variance in hip loading and 5%–87% in knee loading. FNA generally made the largest relative contribution to the variance explained in joint loading across regression models, although its associations varied in direction. Each 10° increase in FNA reduced the first hip compressive and resultant peaks by 0.076 and 0.035 BW, respectively, while hip shear force showed the largest increase, averaging 0.055 BW across both peaks. Most knee loads also increased, by up to 0.131 BW for the second resultant peak. Associations with ETT were primarily observed at the second peak. Our findings suggest that considering gait adaptations due to torsional alterations is crucial for estimating lower-limb joint loading, and prescribing joint kinematics and external forces while altering lower-limb torsion might lead to overestimation.

## 2. Introduction

Idiopathic torsional deformities (ITD) of the femur and tibia are characterised by increased or decreased axial twist of the bone without a known cause, such as a neurological condition or major trauma (Mackay et al., 2021). These deformities may occur unilaterally or bilaterally and involve increased internal or external torsion of the femur and tibia in isolation or combination, with the most common combined presentation being increased femoral torsion, or femoral neck anteversion (FNA), and external tibial torsion (ETT) (Mackay et al., 2021; Passmore et al., 2018). Studies have shown that paediatric patients with ITD are more likely to trip, have reduced functional capacity and societal participation, and experience pain at the hip and knee (Gruskay et al., 2019; Leblebici et al., 2019; Mackay et al., 2021). They also exhibit an altered gait pattern compared with typically developing children (Alexander et al., 2022; Alexander et al., 2024; Alexander et al., 2020; Bruderer-Hofstetter et al., 2015; Byrnes et al., 2020; Hamid et al., 2022; Mackay et al., 2021; Passmore et al., 2018), including differences in hip rotation and foot-progression angle. Similar kinematic adaptations have also been observed in healthy, asymptomatic adults with a broad range of femoral torsion values (De Pieri et al., 2021).

Musculoskeletal modelling and simulation have been applied to non-invasively estimate joint loading, also referred to as joint contact forces, in the presence of anatomical differences, which could help support and guide surgical decision-making for patients with ITD. Existing studies of joint loading have predominantly focused on abnormal FNA, while the impact of excessive ETT on joint loading has not been studied to the best of the authors’ knowledge, only the influence on muscle actions (Hicks et al., 2007; Schwartz and Lakin, 2003). Two methodologies have been used to estimate joint loading in ITD. The first approach prescribes identical gait kinematics and external forces, healthy and pathological depending on the study, across multiple simulations while altering the model’s femoral torsion. These studies consistently found that increasing FNA increases hip and knee joint contact forces (Heller et al., 2001; Kainz et al., 2020; Kainz et al., 2023; Modenese et al., 2021; Passmore et al., 2018; Roth et al., 2021; Shepherd et al., 2022). However, this methodology prevents the emergence of gait adaptations in response to altered femoral torsion. This is problematic because greater internal hip rotation and a more internally directed foot-progression angle are well documented in individuals with increased FNA (Alexander et al., 2022; Alexander et al., 2024; Bruderer-Hofstetter et al., 2015; Hamid et al., 2022; Mackay et al., 2021; Passmore et al., 2018), and this adaptation is hypothesised to improve the hip abductor moment arms (Arnold et al., 1997), which may also influence joint loading. The second approach combines motion capture data with musculoskeletal models incorporating participant-specific femoral torsion and standard inverse-based simulations (Alexander et al., 2022; De Pieri et al., 2021). Findings from these studies have been inconsistent, with increased FNA linked to higher or lower joint loading depending on the population studied (adolescents with increased FNA versus healthy adults with heterogeneous FNA).

A predictive simulation approach may overcome the limitations of previous studies by enabling gait dynamics to emerge in response to independent and combined alterations in FNA and ETT within the same baseline musculoskeletal model. Therefore, the aim of the current study was to use predictive simulation to investigate how isolated and combined changes in FNA and ETT affect lower-limb joint loading when considering kinematic adaptations.

## 3. Methods

### 3.1 Musculoskeletal model

For this study, we used the musculoskeletal model provided by Falisse et al. (2022), which was scaled to match a healthy female adult (mass: 62 kg; height: 1.70 m), as our baseline. The model represented the human skeleton as a multibody system with 20 rigid segments and 31 degrees-of-freedom (DoFs) (Hamner et al., 2010). The lower-limb and trunk DoFs were actuated by 92 muscle-tendon units, whereas the upper-limb DoFs were actuated by 8 coordinate actuators (150 Nm limit) with activation dynamics. Each muscle-tendon unit was modelled using a three-element Hill-type model with contraction (De Groote et al., 2016) and activation dynamics (De Groote et al., 2009). Further model details can be found in prior studies (Falisse et al., 2022; Falisse et al., 2019b). We used polynomial functions with a reduced set of coefficients to describe the lengths, velocities, and moment arms of the muscle-tendon units as functions of the model’s generalised coordinates and velocities (Falisse et al., 2019b; Harba et al., 2026).

### 3.2 Modelling torsional deformities

We unilaterally altered the model’s femoral and tibial torsion using the open-source bone deformation toolbox developed by Modenese et al. (2021). This toolbox applies a user-defined linear rotational profile along a bone’s longitudinal axis, generating a modified skeletal anatomy while adjusting the relevant muscle attachments. FNA and ETT in the baseline model were estimated from the bone geometries as 14° and 29°, respectively. We used the deformation toolbox to generate models with independently varied FNA angles of 1°, 28°, 38°, and 48° and ETT angles of 12°, 20°, 40°, and 53° (see extremes in Figure 1) that were based on the ranges from cadaveric measurements (Strecker et al., 1997). These ranges encompassed previously used thresholds for classifying excessive FNA (30°) (Alexander et al., 2022) and ETT (41°) (Alexander et al., 2020). We also generated models with combined femoral and tibial torsion variations, resulting in a total of 25 models.

**Figure 1.**
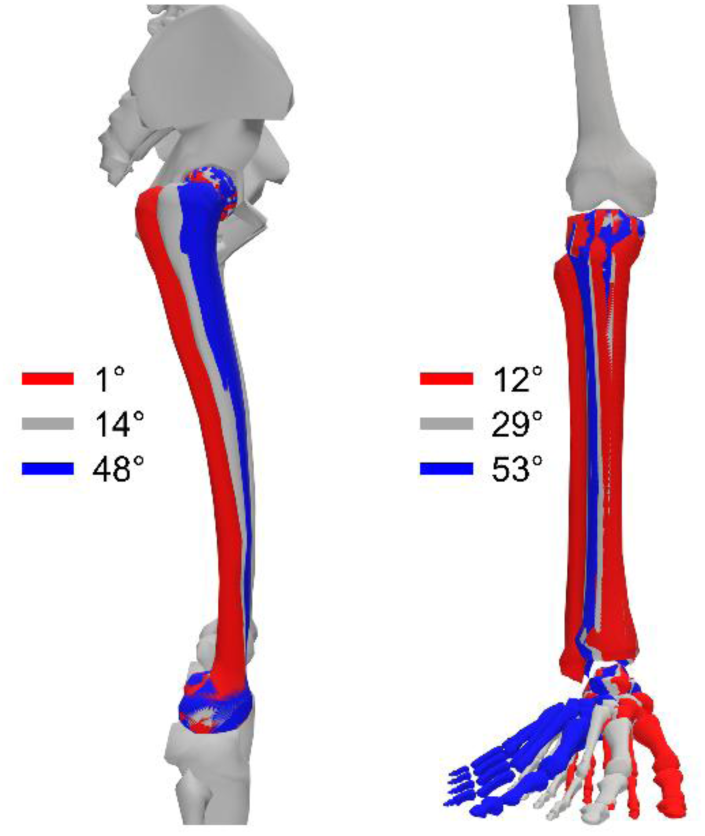
Femoral neck anteversion (left) and external tibial torsion (right) are illustrated for the model’s reference anatomy (grey) and for the lower (red) and upper (blue) extremes of the isolated variations.

### 3.3 Predictive simulations

We used PredSim (D’Hondt et al., 2024b; Falisse et al., 2019b) to generate predictive simulations of walking a whole stride. PredSim is an open-source framework that can be used to formulate predictive simulations as optimal control problems, which are solved as nonlinear programming problems using direct collocation, and supports algorithmic differentiation of multibody dynamics (Falisse et al., 2019a). For each model, we predicted the gait pattern that minimized a cost function previously shown to generate realistic simulations of walking without tracking experimental data (Falisse et al., 2022; Falisse et al., 2019b). The cost function did not include a term explicitly minimising joint contact forces. Simulations were constrained by multibody dynamics, muscle contraction and activation dynamics, coordinate actuator activation dynamics, inter-limb distance, stride periodicity, and walking speed (1.33 m/s). We modified the default settings of PredSim by increasing the upper bound on internal hip rotation (from 27° to 45°; however, no simulations reached this limit) to better capture the range reported in the literature (Passmore et al., 2018). We also lowered the upper bound on knee extension to 0° and decreased the lower bound on muscle activations from 0.05 to 0. All simulations were formulated in MATLAB (2022b; MathWorks Inc., Natick, MA, USA) using CasADi (v3.7.2) (Andersson et al., 2018), and solved using IPOPT (Wächter and Biegler, 2005) with 100 mesh intervals and a convergence tolerance of 10^-4^. For each model, we performed simulations with the default data-informed initial guess provided with PredSim and variants created by adding Gaussian noise with a standard deviation of 2.5×10^-3^. The simulation with the lowest overall objective function value was used in further analysis.

### 3.4 Outcomes

We extracted joint angles and foot-progression angle for the altered lower limb to assess whether the simulations captured kinematic features of torsional deformities reported previously. Foot-progression angle was calculated from the rotation matrix of the calcaneus body expressed in the ground frame, as the projected angle between the calcaneus and ground anterior-posterior axes. Positive foot-progression angles indicate an internally rotated foot. We then used OpenSim’s Joint Reaction Analysis tool (Seth et al., 2018; Steele et al., 2012) to calculate hip, knee and ankle joint contact forces using the models with their geometry-based representation of muscle-tendon units. For each joint, we calculated the axial compressive and transverse shear contact forces, acting on the child segment and expressed in the child segment frame, as well as the resultant. Compressive force was defined as the component parallel to the bone longitudinal axis, while shear force was calculated as the resultant of the perpendicular anterior-posterior and medial-lateral components. Joint contact forces were normalised to body weight (BW), and peak hip and knee contact forces of the altered lower limb were extracted. We focused our analysis on the hip and knee contact forces as individuals with ITD most commonly report pain at these joints (Mackay et al., 2021). Ankle contact forces are provided in the Supplementary Material (Figures S1-S3). To quantify the sensitivity of joint loading to lower-limb torsion, separate additive multiple linear regression models were fitted for the first and second peaks of hip and knee compressive, shear, and resultant forces using MATLAB (2026a; MathWorks Inc., Natick, MA, USA). Regression models including a femoral torsion × tibial torsion interaction term were also examined, but interaction effects were found negligible (maximum coefficient magnitude = 2e-04) and provided only a small increase in explained variance (mean adjusted *R*^2^ = 0.03). Consequently, additive regression models were used to assess the independent effects of femoral and tibial torsion.

## 4. Results

### 4.1 Kinematics

The largest kinematic adaptations to variations in lower-limb torsion were observed in the transverse plane, particularly in hip internal-external rotation (Figure 2) and foot-progression angle (Figure 3). Independently increasing FNA resulted in greater hip internal rotation and a more internally rotated foot-progression angle. Increasing ETT independently also led to greater hip internal rotation, but a more externally rotated foot-progression angle. In the simulations combining torsional changes, hip internal rotation increased with both increased FNA and increased ETT, as expected. In contrast, foot-progression angle depended on the relative contribution of each torsional change. Increasing FNA while reducing ETT produced a more internally rotated foot-progression angle, whereas increasing both FNA and ETT led to minimal change, with the foot remaining externally rotated. Comprehensive joint kinematic waveforms normalised to the gait cycle for the complete set of isolated and combined conditions are provided in the Supplementary Material (Figures S4-S10).

**Figure 2.**
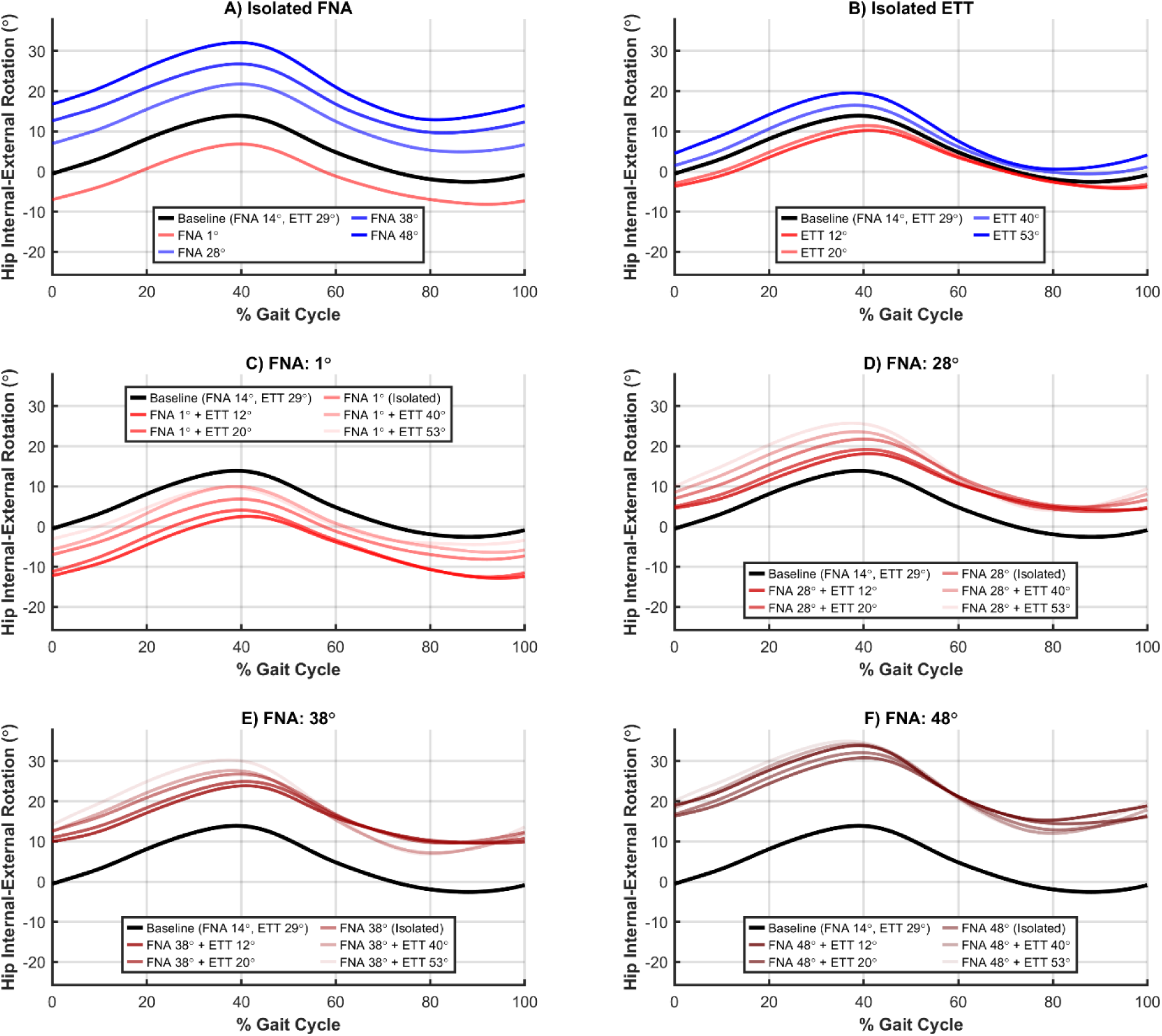
Hip internal-external rotation (positive values indicate internal rotation) over the gait cycle for isolated and combined femoral neck anteversion (FNA) and external tibial torsion (ETT) conditions. The first row shows isolated FNA (A) and ETT (B) conditions. The bottom two rows (C-F) show fixed FNA angles with varied ETT angles.

**Figure 3.**
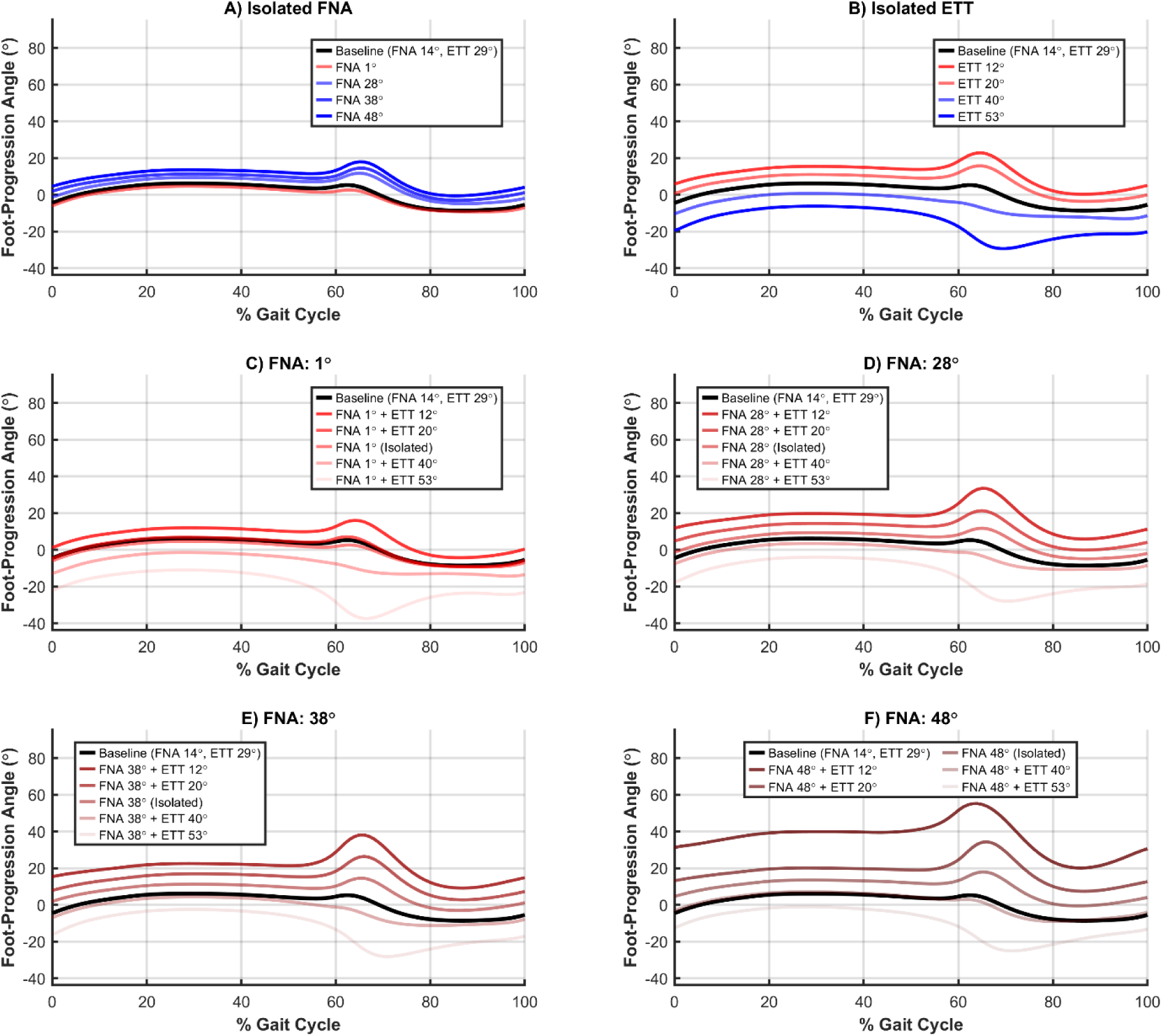
Foot-progression angle (positive values indicate internal foot rotation) over the gait cycle for isolated and combined femoral neck anteversion (FNA) and external tibial torsion (ETT) conditions. The first row shows isolated FNA (A) and ETT (B) conditions. The bottom two rows (C-F) show fixed FNA angles with varied ETT angles.

### 4.2 Joint contact forces

Hip contact force waveforms normalised to the gait cycle are presented for the isolated FNA and ETT conditions in Figure 4, with the complete set of isolated and combined conditions provided in the Supplementary Material (Figures S11-S13). Across the first and second peaks, the regression models explained 23% to 86% of the variation in the magnitude of hip compressive, shear, and resultant forces (Table 1, Figure 5). The strongest regression models were found for the first peaks of compressive and shear force, for which FNA and ETT together accounted for 85% and 86% of the variance, respectively. FNA generally accounted for a larger proportion of the explained variation across hip joint loading outcomes (Figure 6) and led to greater changes in joint loading. Each 10° increase in FNA was associated with reductions of 0.076 and 0.035 BW in the first compressive and resultant force peaks, respectively. Conversely, the same increase in FNA was associated with increases of 0.064 and 0.046 BW in the first and second shear force peaks, respectively, and a 0.043 BW increase in the second resultant force peak. Independent associations with ETT were limited to the second peak, with each 10° increase associated with reductions of 0.037, 0.031, and 0.048 BW in compressive, shear, and resultant force, respectively.

**Figure 4.**
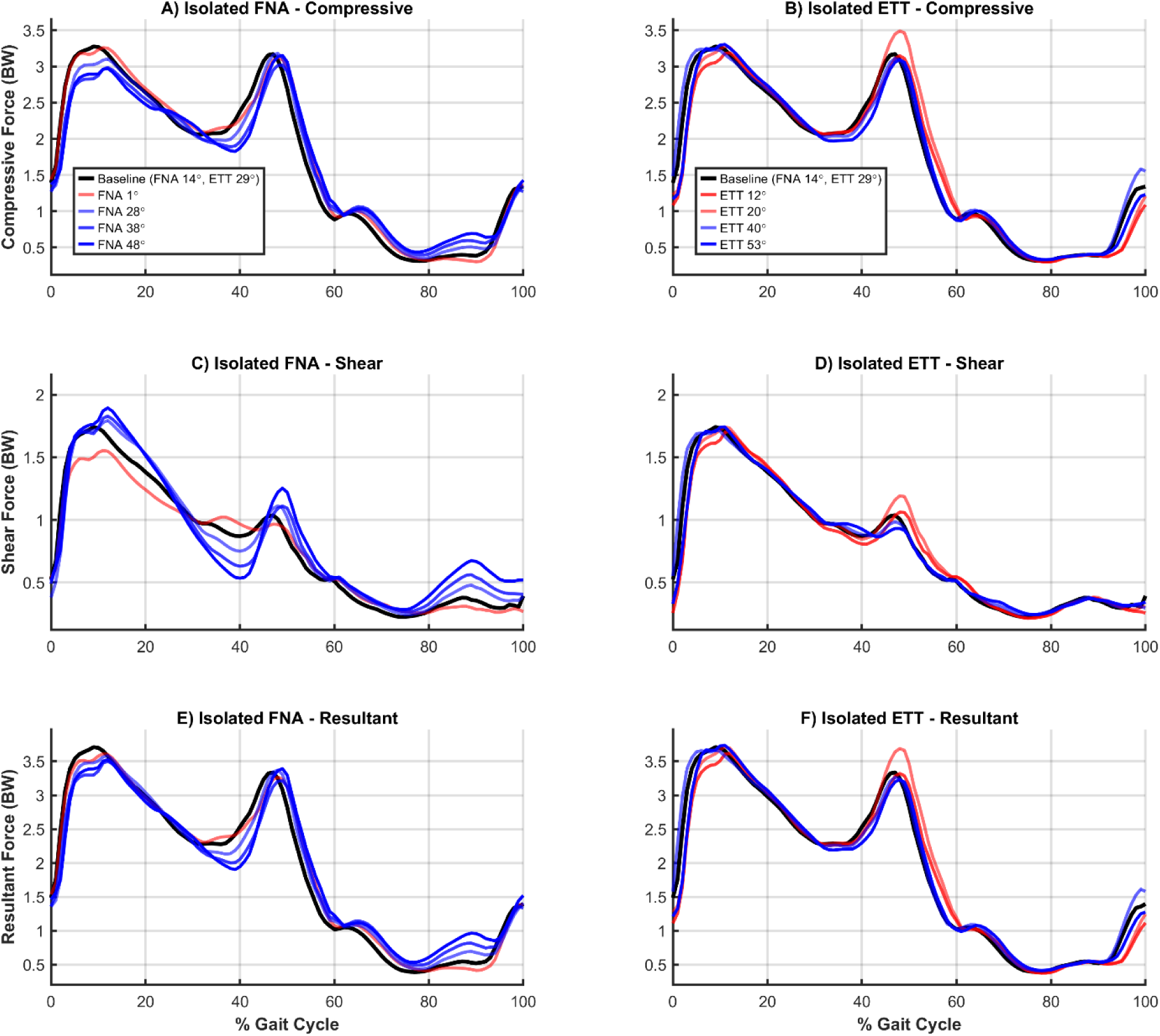
Hip contact forces normalised to body weight (BW) over the gait cycle for isolated femoral neck anteversion (FNA) and external tibial torsion (ETT) conditions. Columns distinguish independently varying either FNA or ETT; rows represent the compressive (A, B), shear (C, D), and resultant (E, F) force components.

**Table 1.** Multiple linear regression results for hip and knee joint loading. Overall model and predictor statistics for the first and second peaks across all force components are provided. 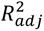: adjusted *R*^2^; CI: confidence interval; FNA: femoral neck anteversion; ETT: external tibial torsion; *B*: unstandardised coefficient (representing the change in body weights per 1° increase in torsion, whereas in-text results scale these coefficients to 10° change to facilitate clinical interpretation).

| Joint & Force Component | Peak | Model $R^2_{adj}$ | FNA $B$<br>(95% CI) | ETT $B$<br>(95% CI) |
| --- | --- | --- | --- | --- |
| Hip Joint |  |  |  |  |
| Compressive | First | 0.85 | -0.0076<br>(-0.0090, -0.0062) | 0.0013<br>(-0.0003, 0.0029) |
|  | Second | 0.23 | 0.0025<br>(-0.0003, 0.0054) | -0.0037<br>(-0.0070, -0.0005) |
| Shear | First | 0.86 | 0.0065<br>(0.0053, 0.0076) | 2.32e-05<br>(-0.0013, 0.0013) |
|  | Second | 0.66 | 0.0047<br>(0.0031, 0.0063) | -0.0032<br>(-0.0050, -0.0013) |
| Resultant | First | 0.42 | -0.0035<br>(-0.0051, -0.0018) | 0.0011<br>(-0.0008, 0.0031) |
|  | Second | 0.38 | 0.0043<br>(0.0012, 0.0073) | -0.0048<br>(-0.0084, -0.0013) |
| Knee Joint |  |  |  |  |
| Compressive | First | 0.58 | 0.0038<br>(0.0025, 0.0051) | 0.0001<br>(-0.0014, 0.0016) |
|  | Second | 0.87 | 0.0129<br>(0.0108, 0.0151) | -0.0029<br>(-0.0054, -0.0004) |
| Shear | First | 0.75 | 0.0071<br>(0.0054, 0.0089) | -0.0015<br>(-0.0035, 0.0006) |
|  | Second | 0.05 | 0.0010<br>(-0.0005, 0.0024) | -0.0009<br>(-0.0027, 0.0008) |
| Resultant | First | 0.66 | 0.0056<br>(0.0039, 0.0072) | -0.0003<br>(-0.0023, 0.0016) |
|  | Second | 0.87 | 0.0131<br>(0.0109, 0.0153) | -0.0032<br>(-0.0057, -0.0007) |

**Figure 5.**
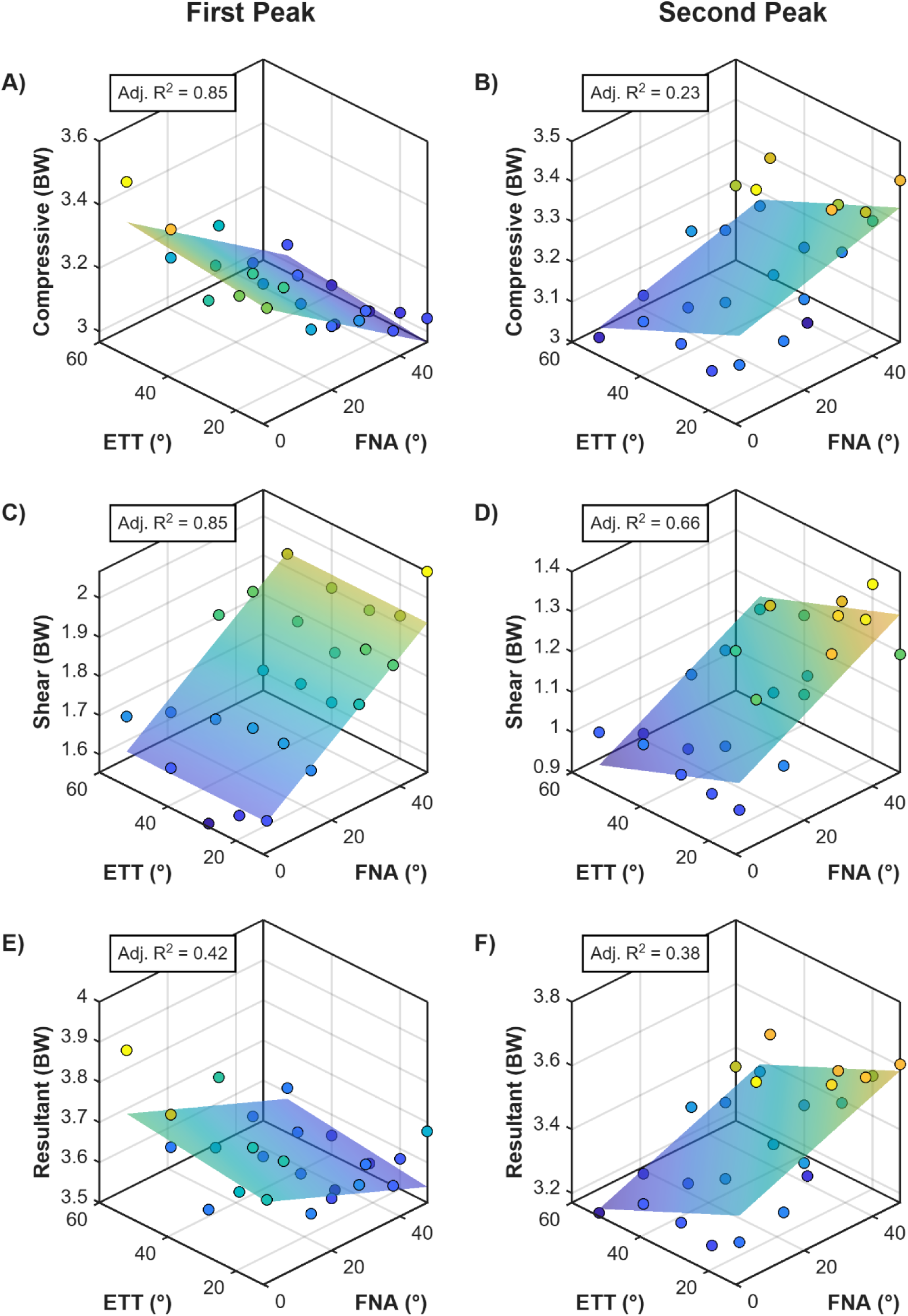
Peak hip contact forces and corresponding linear regression surface fits across variations in femoral neck anteversion (FNA) and external tibial torsion (ETT). Scatter points denote the predictive simulation output, and shaded surfaces represent the regression models. Columns distinguish the first and second peaks; rows represent the compressive (A, B), shear (C, D), and resultant (E, F) force components expressed in body weight (BW). The adjusted R^2^ (Adj. R^2^) for each regression model is included within the respective subplots.

**Figure 6.**
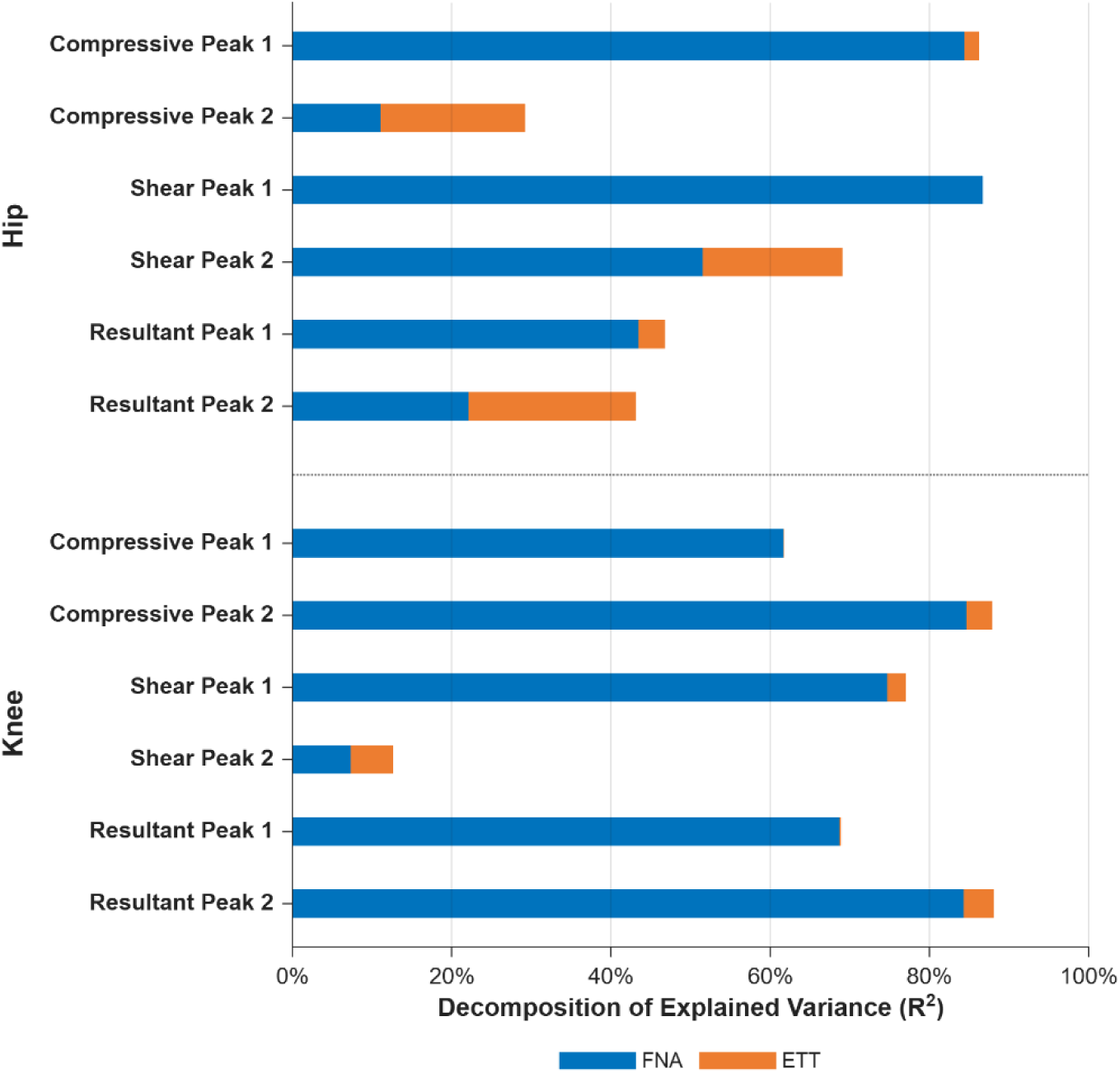
Decomposition of the variance explained by each of the separate multiple linear regression models. Each horizontal bar partitions the model’s total R^2^ into the absolute contributions unique to femoral neck anteversion (FNA) and external tibial torsion (ETT), with the total bar length representing the combined model R^2^.

Knee contact force waveforms normalised to the gait cycle are presented for the isolated FNA and ETT conditions in Figure 7, with the complete set of isolated and combined conditions provided in the Supplementary Material (Figures S14-S16). The regression models explained 5% to 87% of the variation across the first and second peaks in the magnitude of knee compressive, shear, and resultant forces (Table 1, Figure 8). The strongest regression models were found for the second peaks of compressive and resultant force (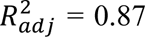 for both), while the weakest model was found for the second shear force peak (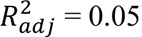). As with hip joint loading, FNA accounted for a greater proportion of the explained variation across knee joint loading (Figure 6) and produced larger changes in force. Each 10° increase in FNA was associated with increases of 0.038 and 0.129 BW in the first and second compressive peaks, respectively, a 0.071 BW increase in the first shear force peak, and increases of 0.056 and 0.131 BW in the first and second resultant force peaks, respectively. Associations with ETT were limited to the second compressive and resultant force peaks, with each 10° associated with reductions of 0.029 and 0.032 BW, respectively.

**Figure 7.**
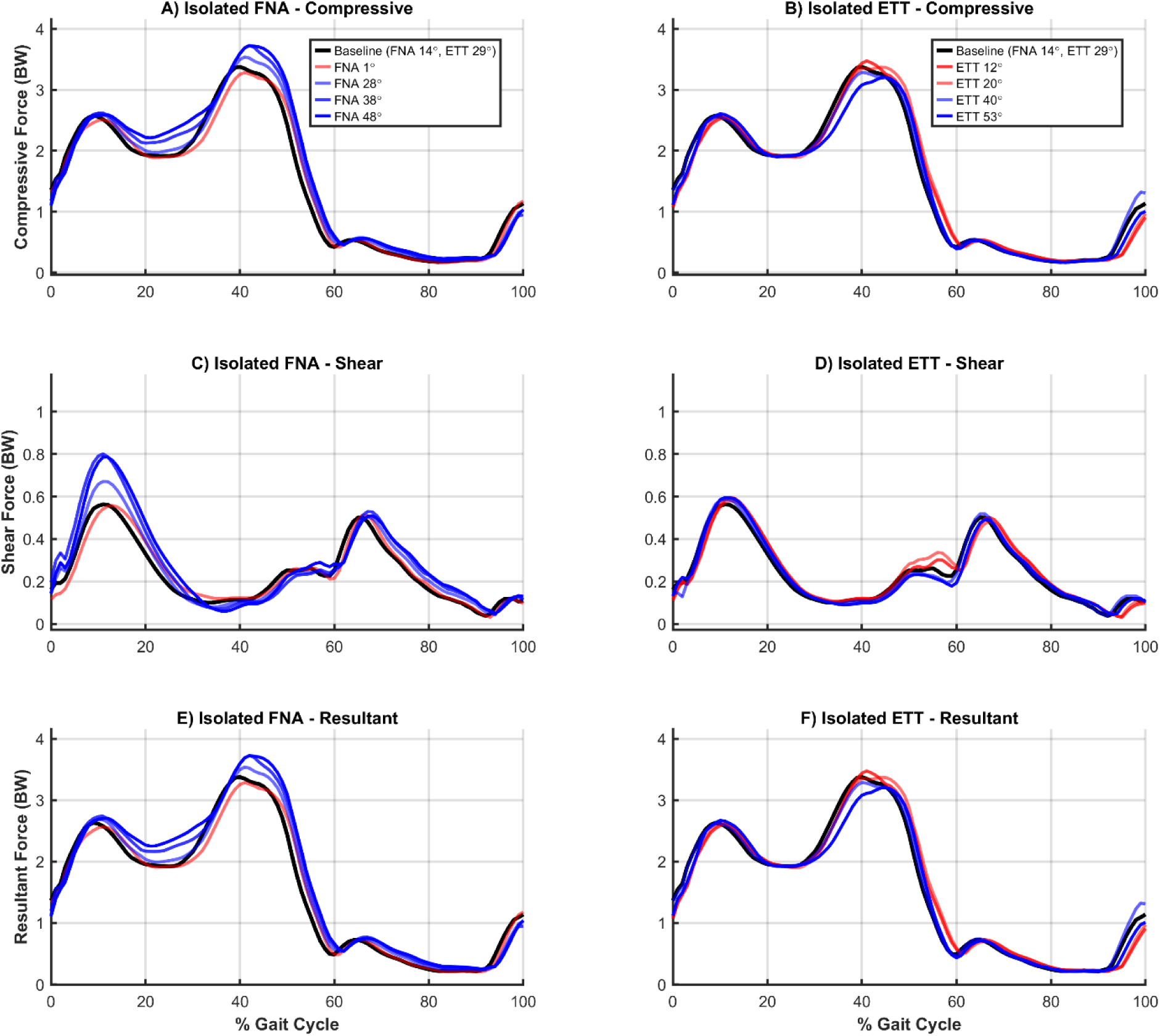
Knee contact forces normalised to body weight (BW) over the gait cycle for isolated femoral neck anteversion (FNA) and external tibial torsion (ETT) conditions. Columns distinguish independently varying either FNA or ETT; rows represent the compressive (A, B), shear (C, D), and resultant (E, F) force components.

**Figure 8.**
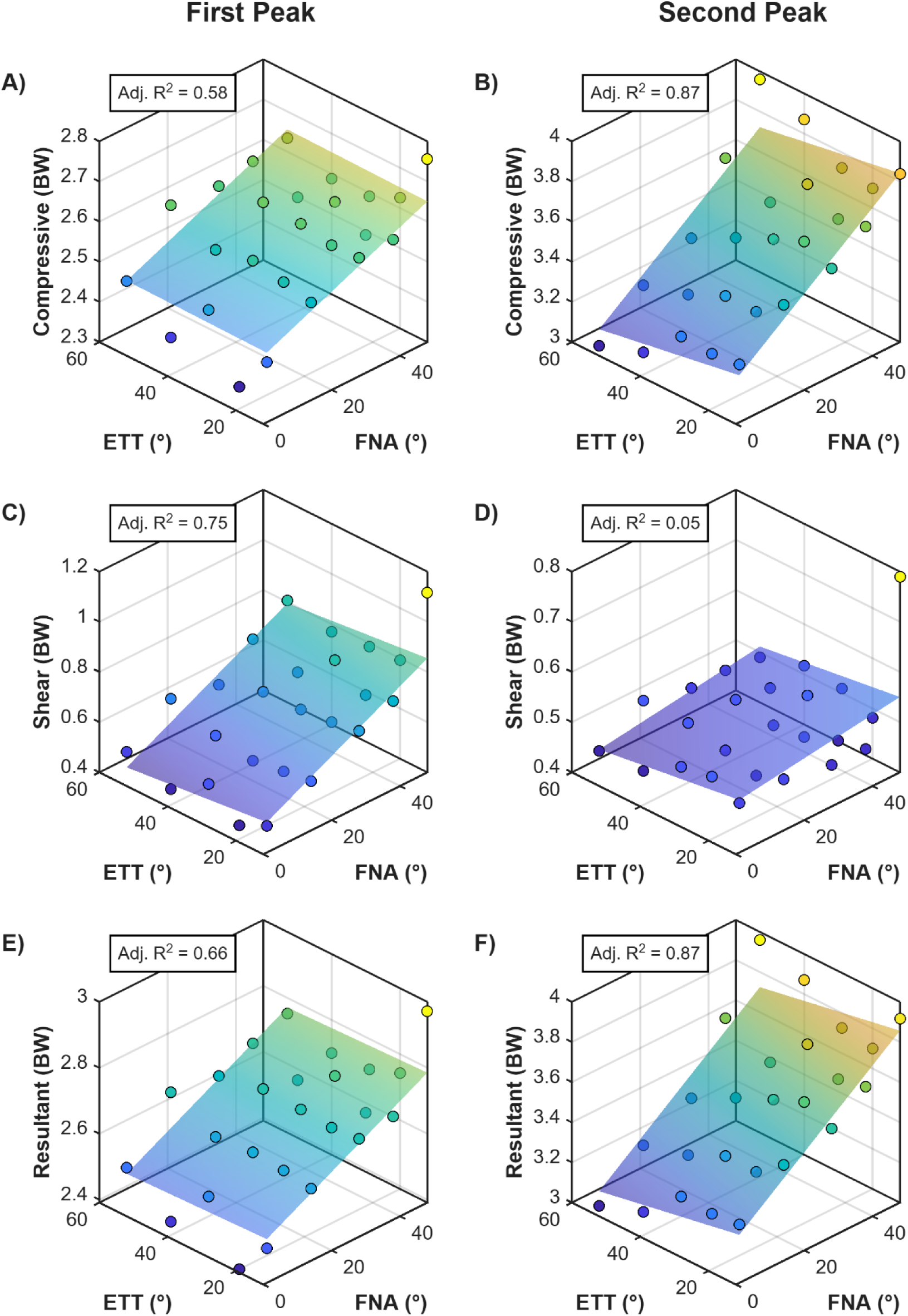
Peak knee contact forces and corresponding linear regression surface fits across variations in femoral neck anteversion (FNA) and external tibial torsion (ETT). Scatter points denote the predictive simulation output, and shaded surfaces represent the regression models. Columns distinguish the first and second peaks; rows represent the compressive (A, B), shear (C, D), and resultant (E, F) force components expressed in body weight (BW). The adjusted R^2^ (Adj. R^2^) for each regression model is included within the respective subplots.

## 5. Discussion

The aim of this study was to investigate how FNA and ETT affect hip and knee joint loading, when kinematic adaptations are enabled by a predictive simulation approach for studying walking with varied lower-limb torsion.

In our simulations we observed that kinematic adaptations occurred with lower-limb torsion. Compared with the baseline model (FNA 14°, ETT 29°), the model with the largest FNA (48°) exhibited an 18° increase in internal hip rotation and a 13° increase in internal foot-progression. The model with highest ETT (53°) showed a 6° increase in internal hip rotation and reached an external foot-progression angle of 29°. The model combining the largest FNA and ETT showed a 21° increase in internal hip rotation and reached an external foot-progression angle of 25°. These kinematic adaptations are consistent with those observed in previous experimental studies on ITD participants (Alexander et al., 2022; Alexander et al., 2024; Alexander et al., 2020; Bruderer-Hofstetter et al., 2015; Byrnes et al., 2020; Hamid et al., 2022; Mackay et al., 2021; Passmore et al., 2018). These adaptations also align with the compensatory mechanism discussed by Arnold et al. (1997) to improve the moment arm of the gluteus medius. Pelvic retraction and increased obliquity, likely compensating for the increased internal hip rotation, were also observed (Figures S8 and S10). These elements give us confidence about the realism of the predicted gait adaptations.

Our simulations also showed that greater FNA modified the direction of hip loading: it reduced the first compressive peak but shear force at both peaks increased (Figure 4), leading to an increased resultant force at the second peak. At the knee, it increased compressive and resultant forces at both peaks and increased shear force at the first peak (Figure 7). Greater ETT was primarily associated with reduced second peak loading at the hip and knee. Changes in joint contact forces were more consistently associated with FNA than ETT, as indicated by the relative share of explained variance (Figure 6), and our discussion therefore focuses on the findings of altered FNA. For most contact force outcomes, the estimated response to combined lower-limb torsional changes largely followed the response associated with FNA, as indicated by its generally larger contribution to the additive regression estimate. The resultant hip contact force at the second peak was the only exception, for which the contribution of ETT was slightly greater.

The findings of our study only partly agree with previous publications that, by increasing FNA without considering kinematics adaptations, have reported larger hip and knee joint contact force components and resultants (Heller et al., 2001; Kainz et al., 2020; Kainz et al., 2023; Modenese et al., 2021; Passmore et al., 2018; Roth et al., 2021; Shepherd et al., 2022). Our predictive approach, however, overcomes the limitation of those prior studies by enabling gait adaptations to emerge from variations in bone torsion. In contrast to those studies, we found a reduction in the first peak compressive (0.076 BW per 10°) and resultant (0.035 BW per 10°) hip contact forces with increased FNA, suggesting that those previous joint contact force estimates are greater, and likely overestimated, because of prescribing the same joint kinematics. However, greater FNA was found to increase the second resultant contact force peak by 0.043 BW per 10°, indicating that its influence on resultant loading differed between peaks. It is worthwhile noting that, while the change in first peak compressive hip force was strongly described by the additive changes in lower-limb torsion (85% of explained variance), the change in resultant hip contact force may also depend on further factors, such as nonlinear changes in gait dynamics, as only moderate proportions of the variance at the first (42%) and second (38%) peaks were accounted for.

To further confirm that differences in hip contact forces reported from previous studies could be explained by the simulation approach, we calculated hip and knee joint loading using the kinematics and external forces from the predictive simulation with baseline anatomy, applied to both the baseline anatomy model and the model with most extreme FNA, following the simulation approach of those previous studies (Figure 9). We found that, with prescribed kinematics and external forces, the model with most extreme FNA resulted in compressive and resultant hip contact forces that on average across both peaks increased by 0.37 and 0.49 BW (10.5% and 13.1%), respectively. When accounting for the kinematic adaptations, the compressive peaks decreased by 0.20 BW (4.9%), while the resultant contact force decreased by 0.18 BW (5.0%) at the first peak and increased by 0.06 BW (1.7%) at the second. Furthermore, the difference in second peak resultant knee contact force was approximately three times as large in comparison to the equivalent predictive simulation (1.11 BW [33.0%] vs. 0.35 BW [10.3%]). These results support the finding that predictive simulation can capture kinematic adaptations occurring for torsional skeletal alterations, and these adaptations have to be considered otherwise the lower-limb joint loading will be overestimated.

**Figure 9.**
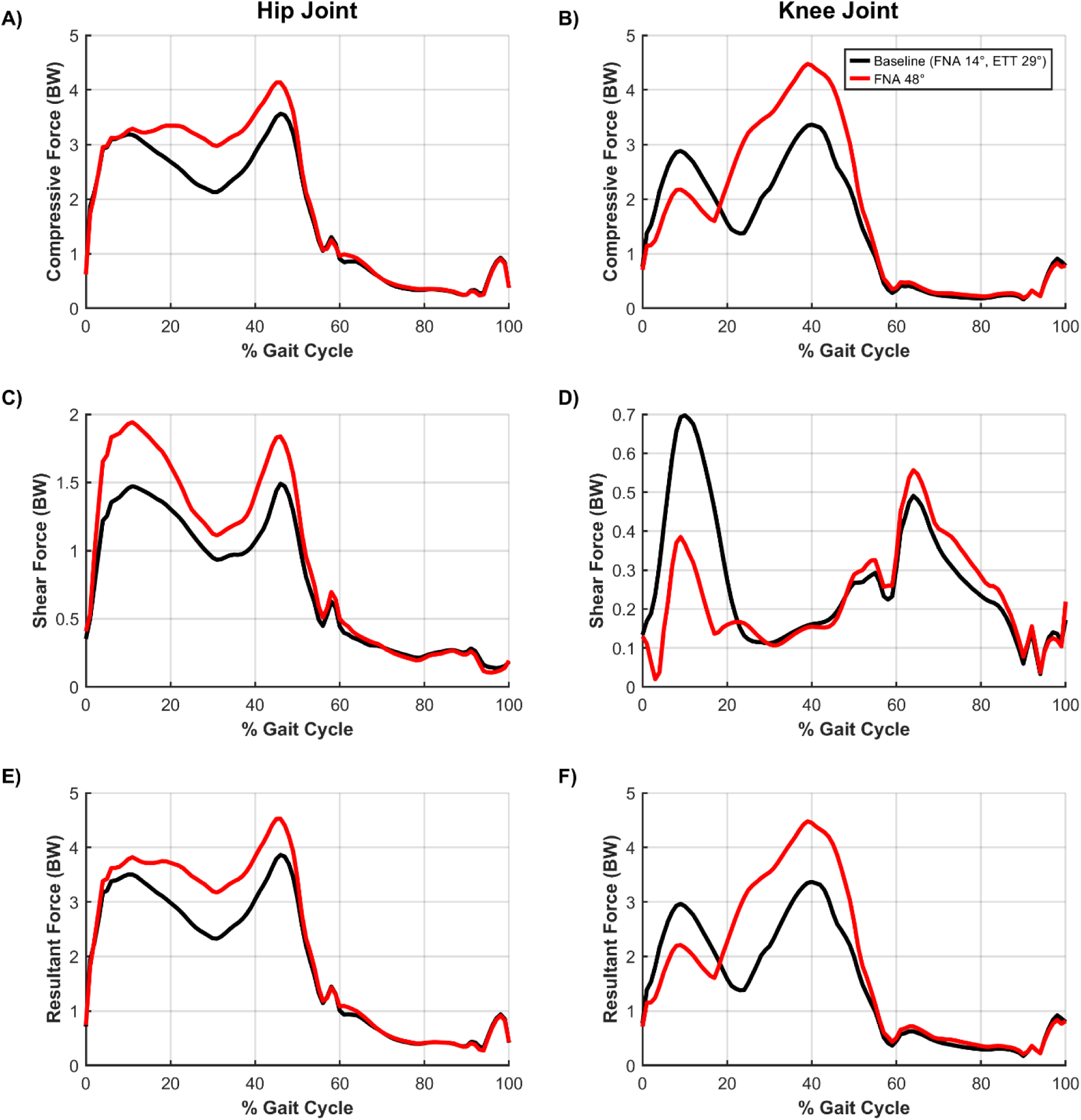
Compressive, shear, and resultant hip and knee joint contact forces, normalised to body weight (BW), over the gait cycle using the kinematics and external forces from the predictive simulation with baseline anatomy, applied to both the baseline model and the model with the most extreme femoral neck anteversion (FNA). Columns distinguish the hip (left) and knee (right) joints. Rows represent the compressive (A, B), shear (C, D), and resultant (E, F) force components. ETT: external tibial torsion.

Our findings were less consistent with those studies using an inverse approach and personalised models to calculate hip and knee joint contact forces. While we found reduced compressive (first peak) and increased shear contact forces at the hip, Alexander et al. (2022) reported lower estimations of both loadings in a paediatric group with elevated FNA compared to typically developing controls. Similarly, they computed decreased knee contact forces, while we observed an increase. De Pieri et al. (2021), on the other hand, reported an association between increased FNA and greater shear hip contact force, in agreement with our findings, but no significant increase in compressive force. Both these studies, however, used a musculoskeletal model with a substantially higher level of discretisation for the lower limb muscles (Carbone et al., 2015; De Pieri et al., 2018) (166 vs 86 muscle bundles in our model), which has been shown to influence joint contact force estimation (Mathai and Gupta, 2019; Weinhandl and Bennett, 2019). It is worth noting, however, that the compressive and shear hip contact forces of Alexander et al. (2022) and De Pieri et al. (2021) are not consistent with each other despite using the same underlying musculoskeletal model. Differences in individual gait patterns and walking speed, which affect both the magnitude and directionality of contact forces (De Pieri et al., 2019; Modenese and Phillips, 2012), as well as differences in cohort ages, particularly the need for Alexander et al. (2022) to scale an adult musculoskeletal model to a paediatric cohort, likely explain these inconsistencies. In contrast, our predictive simulation approach enabled direct control over walking speed and musculoskeletal anatomy while allowing gait to adapt to the imposed torsional changes. Nevertheless, neither of those studies reported the substantial increases in joint loading observed in previous studies using fixed kinematic inputs. This supports the conclusion of our predictive simulation study: neglecting kinematic adaptations may overestimate the effect of torsional deformities on joint loading.

This study presents a few limitations. Firstly, our simulations relied on gradient-based optimisation, which can be sensitive to the initial guess provided and finding local rather than global minima (Schutte et al., 2005). To mitigate this, we varied the initial guess for the design variables when performing the simulations and retained the optimal solution for each varied model with the lowest objective function value. However, due to redundancy in the neuromuscular system, a feasible set of near-optimal solutions likely exists. Within this set, slight variations in muscle recruitment could yield near identical objective function values but result in different joint contact forces, meaning the solution with the lowest objective function value does not necessarily represent the movement strategy with the lowest joint loading. Secondly, our simulations did not fully capture the ankle dorsiflexion-plantarflexion range of motion observed during gait analysis of inverse simulations, typically between approximately 10° dorsiflexion to 20° plantarflexion (Modenese et al., 2018), but were limited to a smaller range (between 15° dorsiflexion and 7° plantarflexion, Figure S4). This limitation has also been identified in previous predictive simulation studies (D’Hondt et al., 2024a; Falisse et al., 2022), which has prompted the development of a new foot model to improve ankle kinematics in predictive simulations (D’Hondt et al., 2024a). However, the improvements were modest for the additional complexity and parameters introduced, and we therefore used the nominal two-segment foot model included in the musculoskeletal model.

## 6. Conclusion

In conclusion, our predictive simulations captured the kinematic adaptations for hip rotation and foot-progression angle reported by experimental studies in individuals with ITD. Moreover, they showed that FNA generally led to greater changes in hip and knee joint loading than ETT. By capturing the kinematic changes across variations in lower-limb torsion, we demonstrated that joint loading estimates differ in both magnitude and direction when compared to altering the model’s FNA while prescribing the joint kinematics and external forces, as commonly done in previous studies. This highlights that joint loading in the presence of torsional deformities cannot reliably be estimated without considering gait adaptations. Future work will focus on clarifying the relationship between torsional deformities and joint loading in paediatric populations aiming to eventually assist with the clinical-decision making surrounding corrective surgery to treat torsional deformities.

## Acknowledgements

This work was supported by the MRFF-funded Motion Connect project (MRFFRDII000028). This research was produced in whole or part by UNSW Sydney researchers and is subject to the UNSW Intellectual property policy. For the purposes of Open Access, the author has applied a Creative Commons Attribution CC-BY license to any Author Accepted Manuscript (AAM) version arising from this submission.

## Data Statement

The data that support the findings of this study will be made available on SimTK (simtk.org).

## Declaration of generative AI and AI-assisted technologies in the manuscription preparation process

During the preparation of this work the author(s) used Microsoft 365 Copilot and Google Gemini to improve the readability of this manuscript. After using these tools/services, the author(s) reviewed and edited the content as needed and take(s) full responsibility for the content of the published article.

