## Supplementary Material for "Joint loading in the presence of torsional deformities is overestimated unless gait adaptations are considered: a predictive simulation approach"

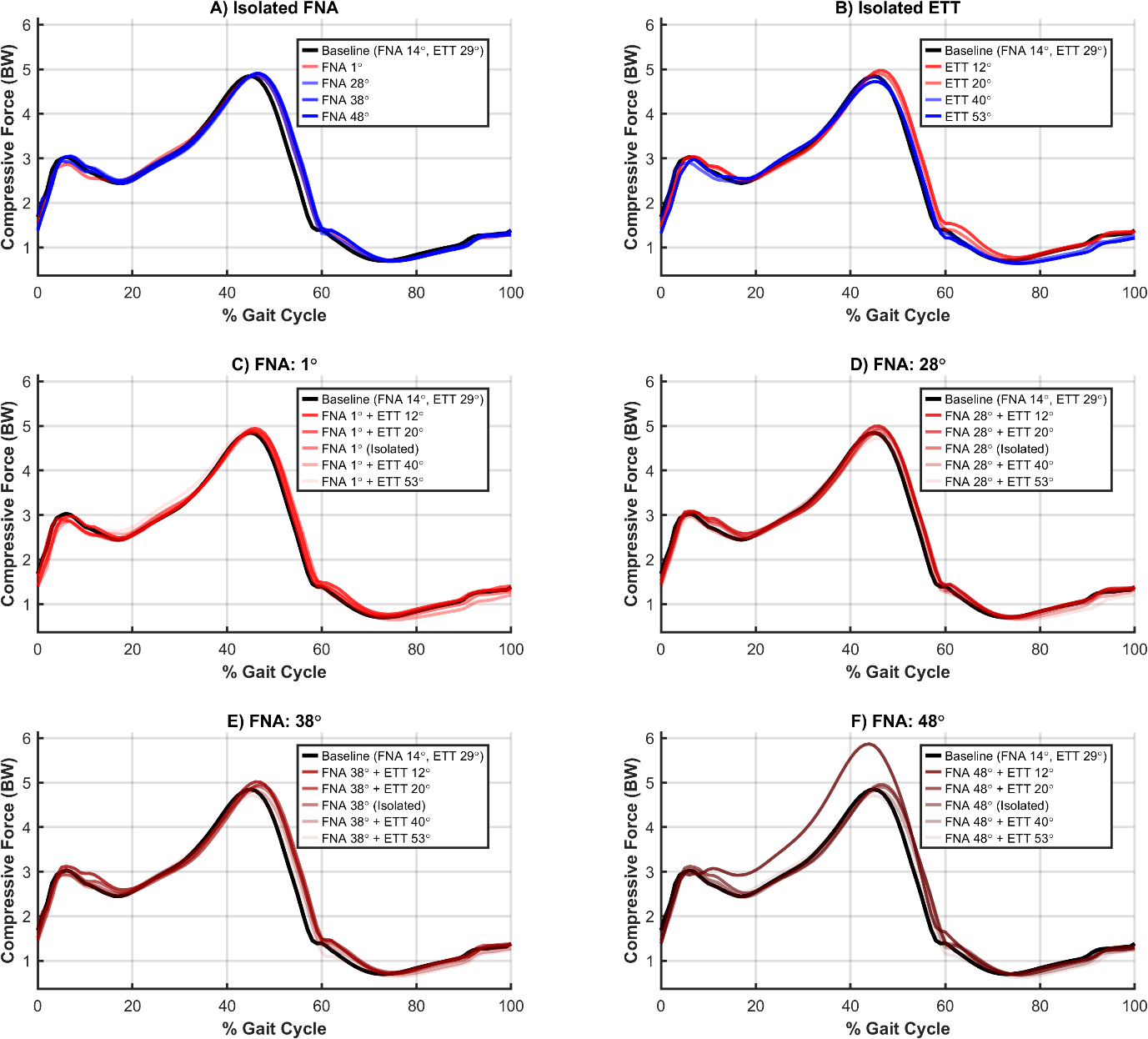

**Supplementary Figure 1** Compressive ankle joint contact force normalised to body weight (BW) over the gait cycle for isolated and combined femoral neck anteversion (FNA) and external tibial torsion (ETT) conditions. The first row shows isolated FNA (A) and ETT (B) conditions. The bottom two rows (C-F) show fixed FNA angles with varied ETT angles.

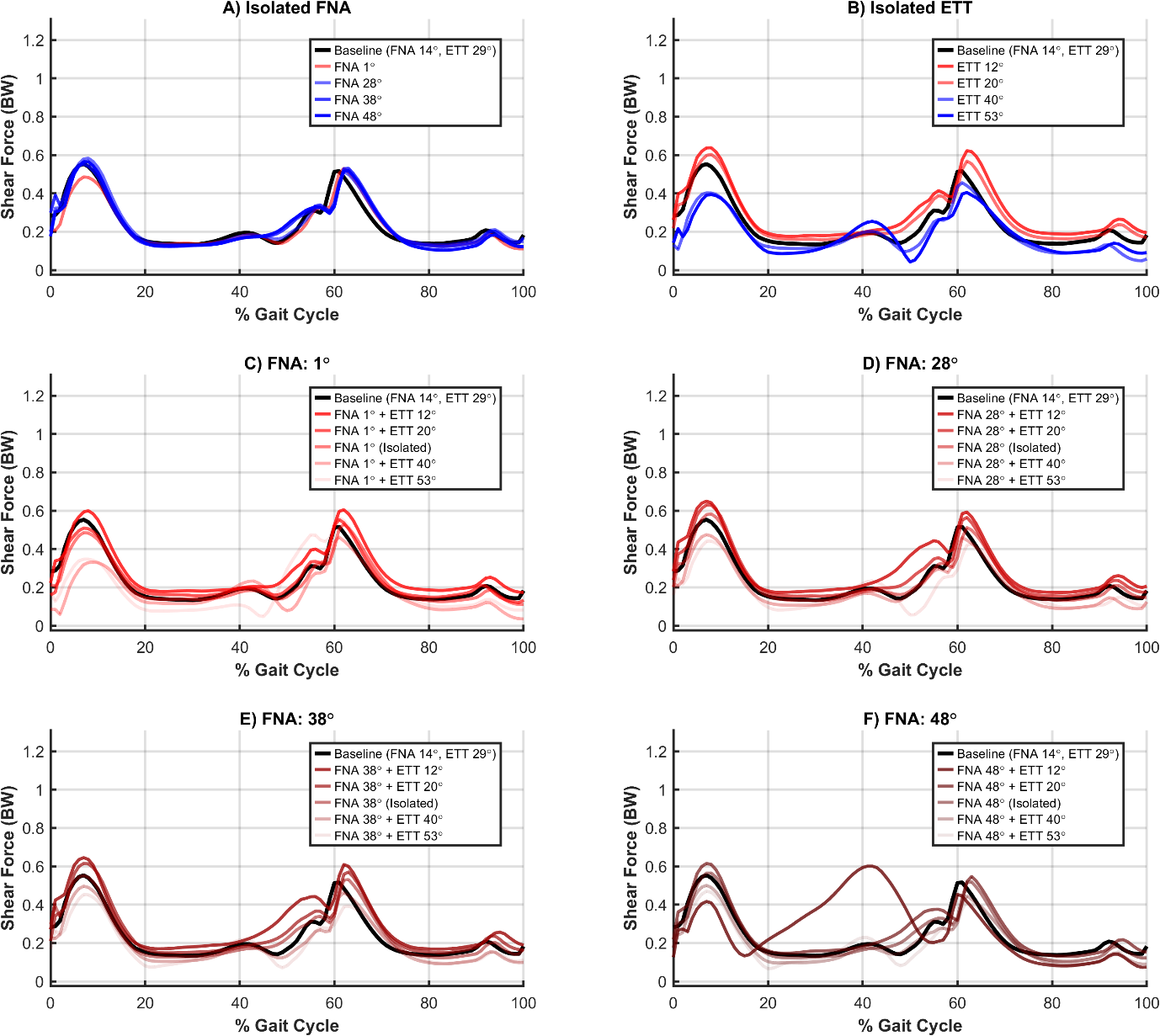

**Supplementary Figure 2** Shear ankle joint contact force normalised to body weight (BW) over the gait cycle for isolated and combined femoral neck anteversion (FNA) and external tibial torsion (ETT) conditions. The first row shows isolated FNA (A) and ETT (B) conditions. The bottom two rows (C-F) show fixed FNA angles with varied ETT angles.

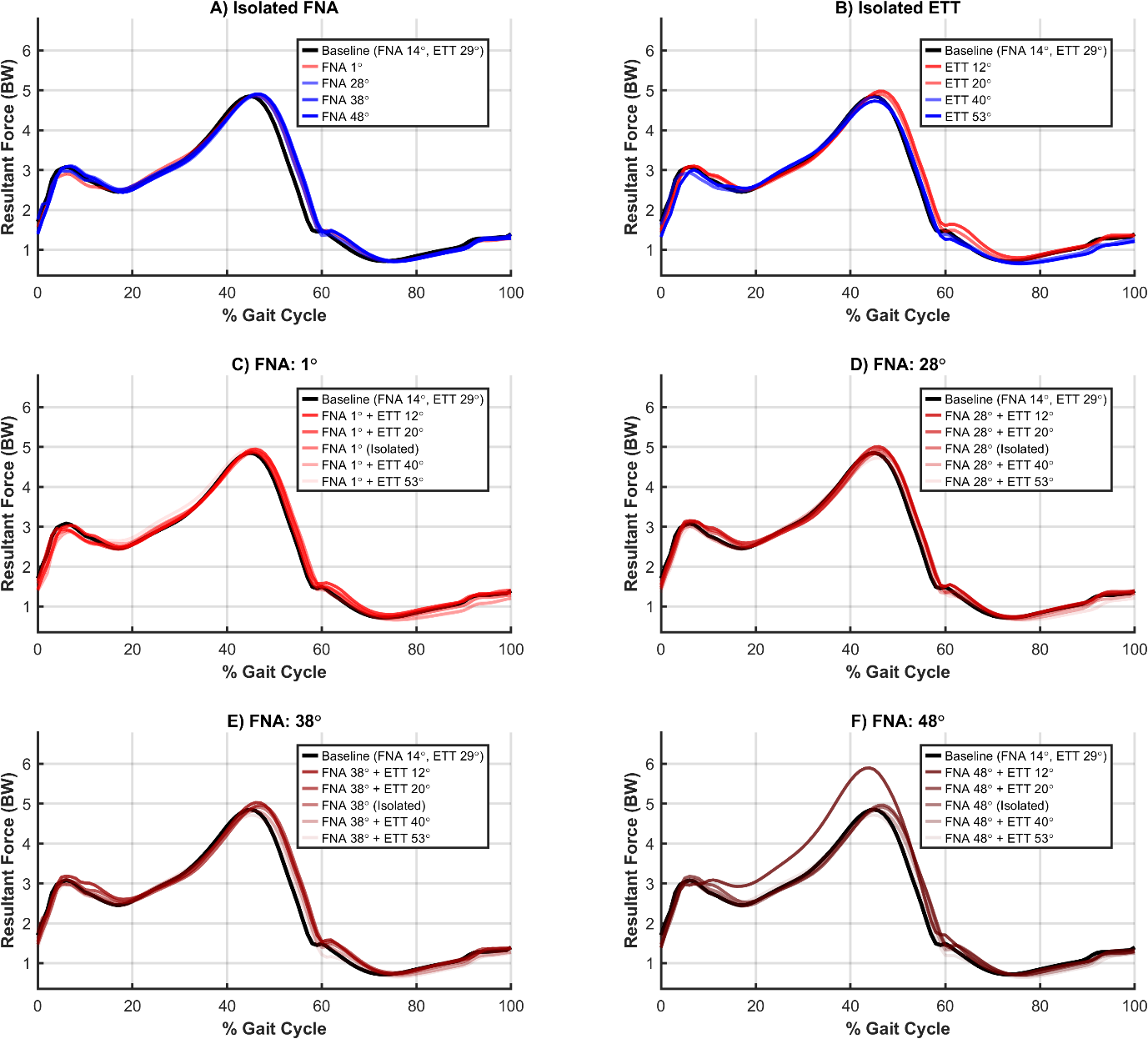

**Supplementary Figure 3** Resultant ankle joint contact force normalised to body weight (BW) over the gait cycle for isolated and combined femoral neck anteversion (FNA) and external tibial torsion (ETT) conditions. The first row shows isolated FNA (A) and ETT (B) conditions. The bottom two rows (C-F) show fixed FNA angles with varied ETT angles.

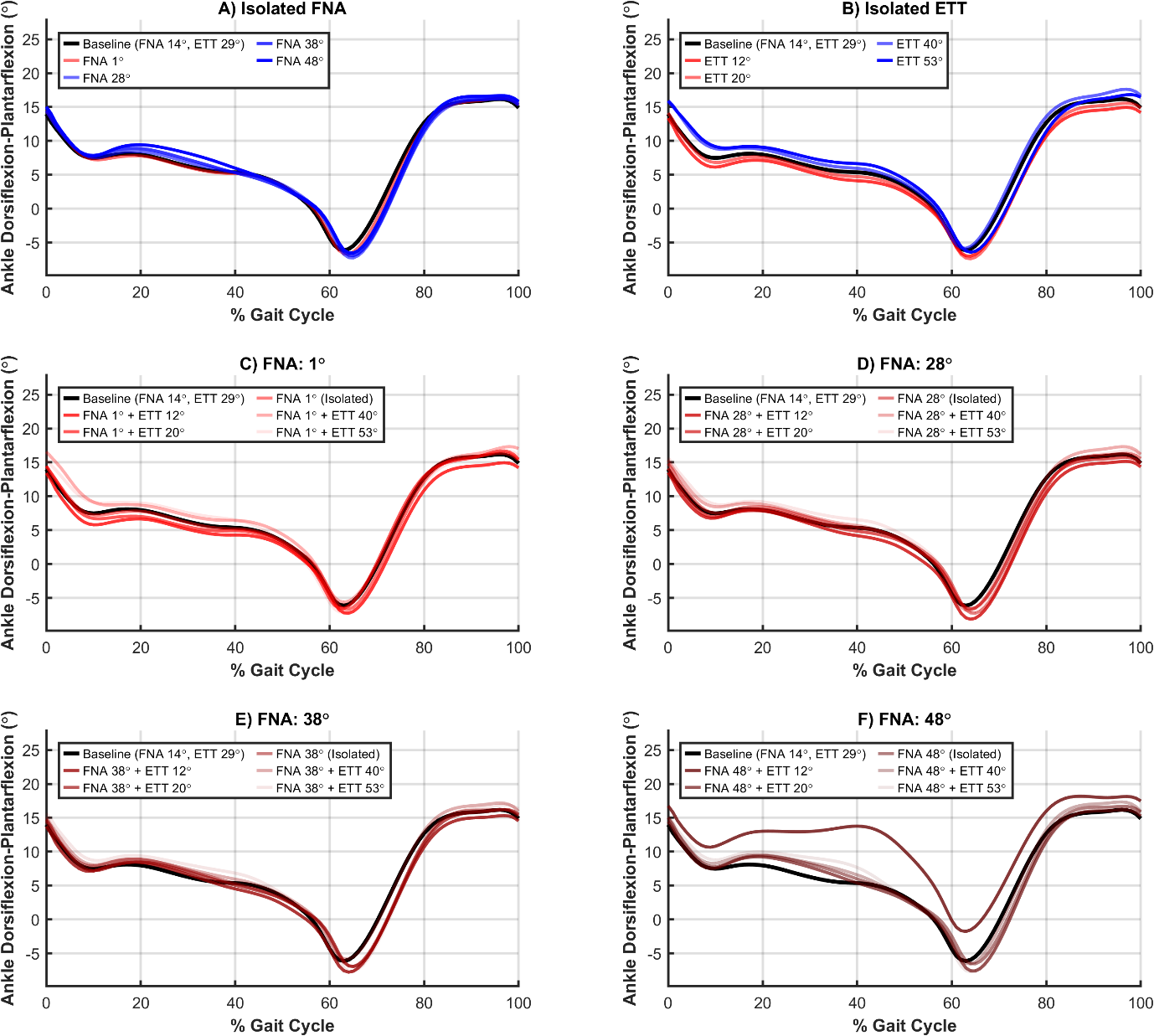

**Supplementary Figure 4** Ankle dorsiflexion-plantarflexion (positive values indicate dorsiflexion) over the gait cycle for isolated and combined femoral neck anteversion (FNA) and external tibial torsion (ETT) conditions. The first row shows isolated FNA (A) and ETT (B) conditions. The bottom two rows (C-F) show fixed FNA angles with varied ETT angles.

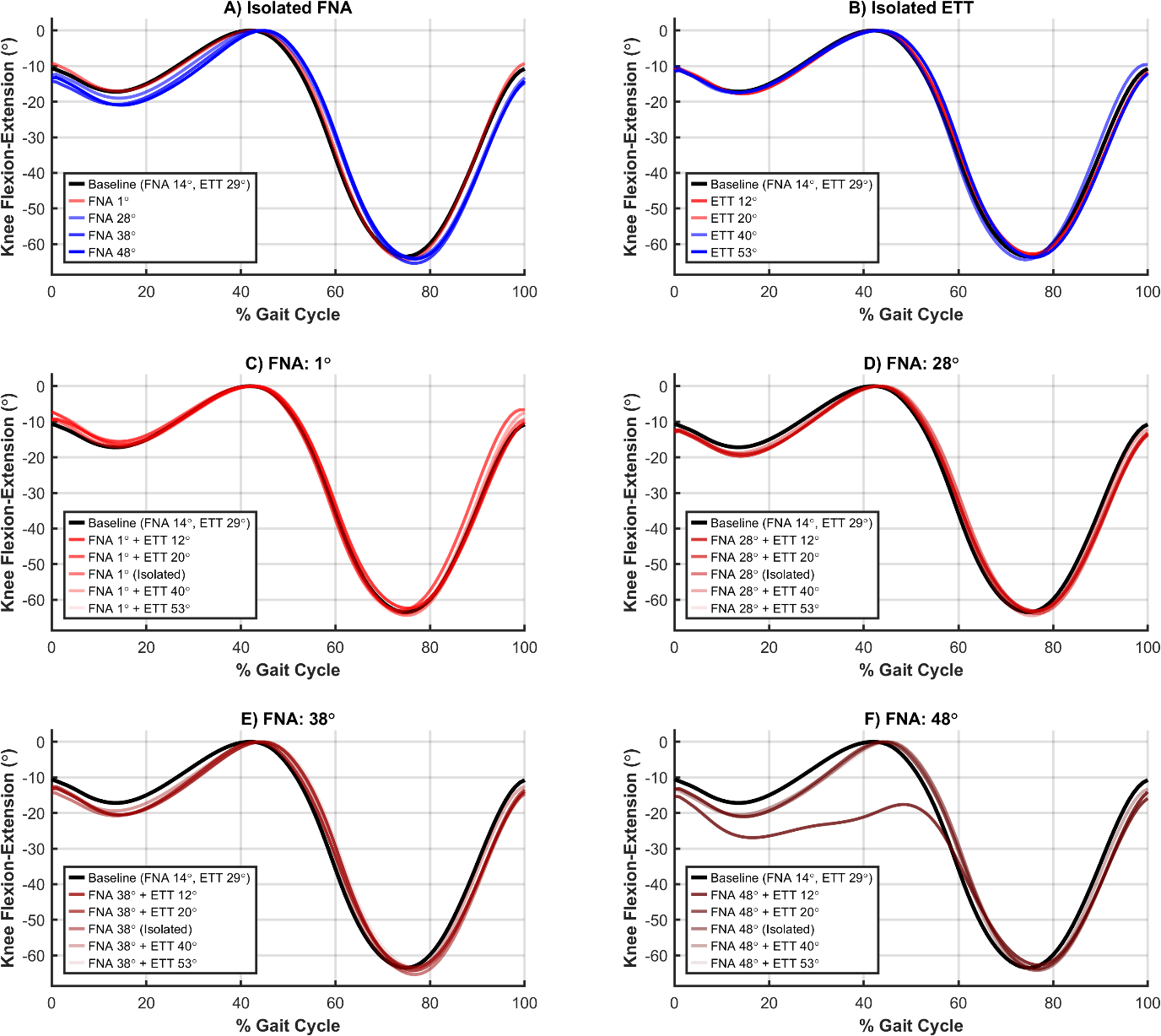

**Supplementary Figure 5** Knee flexion-extension (negative values indicate flexion) over the gait cycle for isolated and combined femoral neck anteversion (FNA) and external tibial torsion (ETT) conditions. The first row shows isolated FNA (A) and ETT (B) conditions. The bottom two rows (C-F) show fixed FNA angles with varied ETT angles.

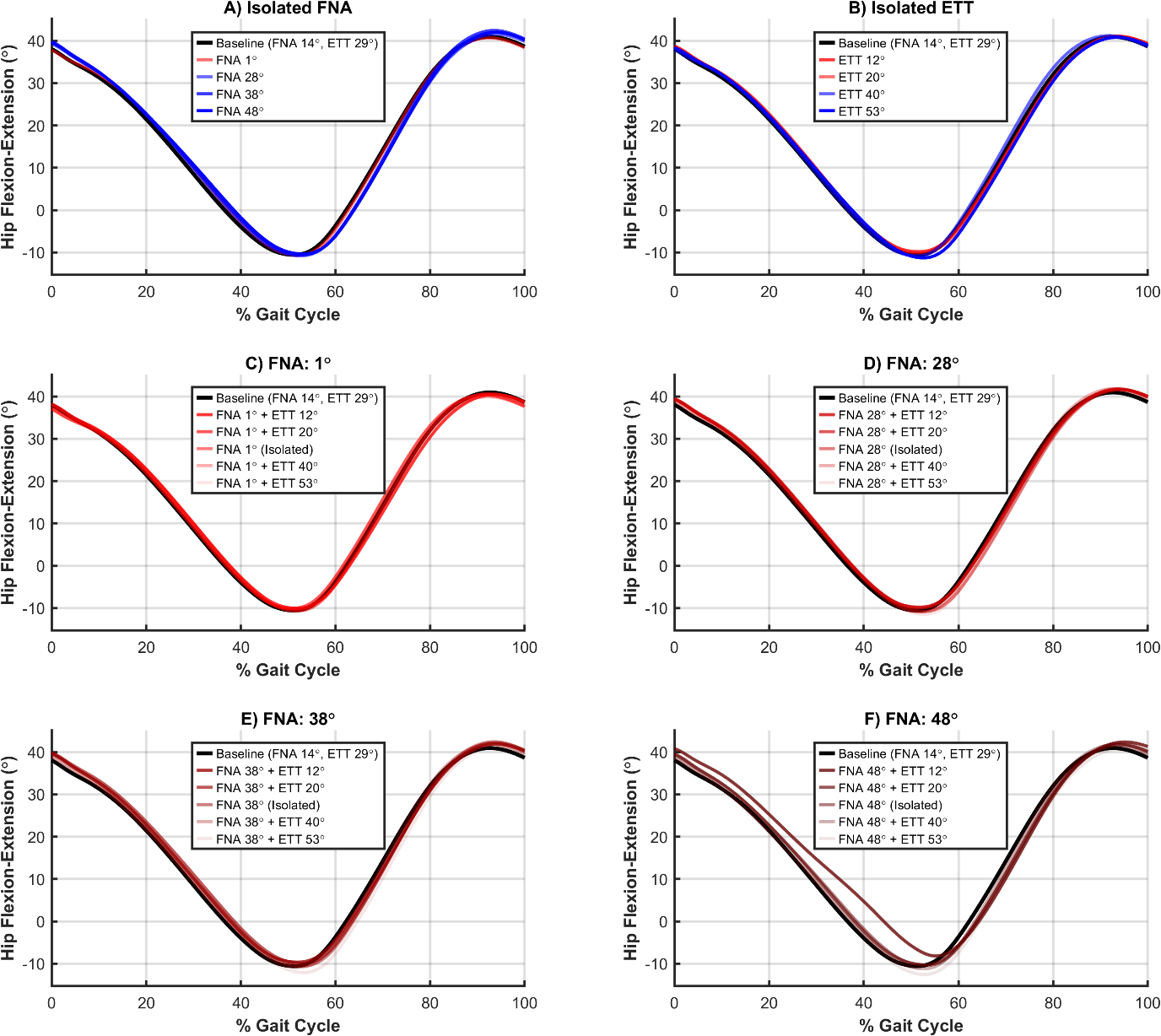

**Supplementary Figure 6** Hip flexion-extension (positive values indicate flexion) over the gait cycle for isolated and combined femoral neck anteversion (FNA) and external tibial torsion (ETT) conditions. The first row shows isolated FNA (A) and ETT (B) conditions. The bottom two rows (C-F) show fixed FNA angles with varied ETT angles.

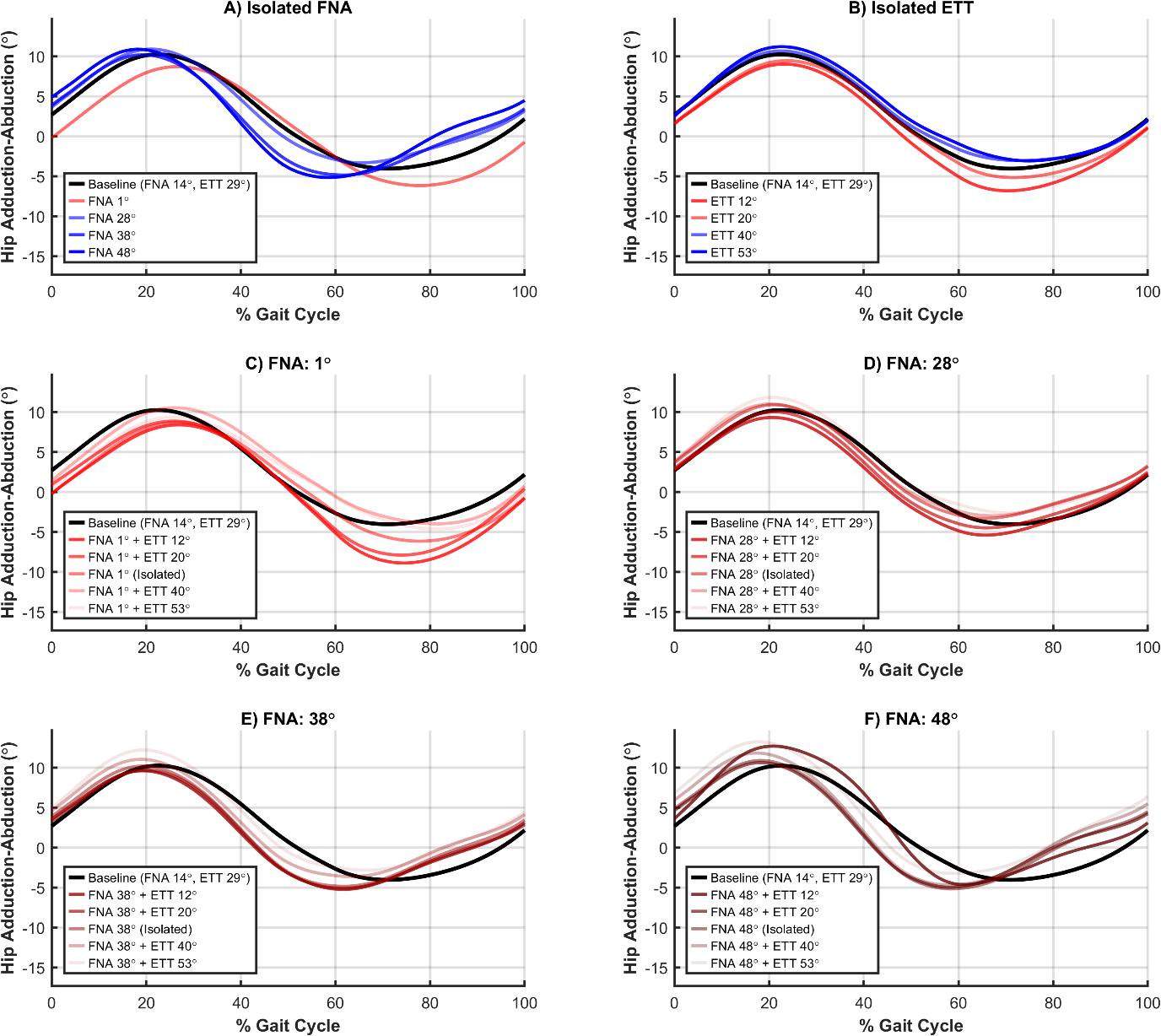

**Supplementary Figure 7** Hip adduction-abduction (positive values indicate adduction) over the gait cycle for isolated and combined femoral neck anteversion (FNA) and external tibial torsion (ETT) conditions. The first row shows isolated FNA (A) and ETT (B) conditions. The bottom two rows (C-F) show fixed FNA angles with varied ETT angles.

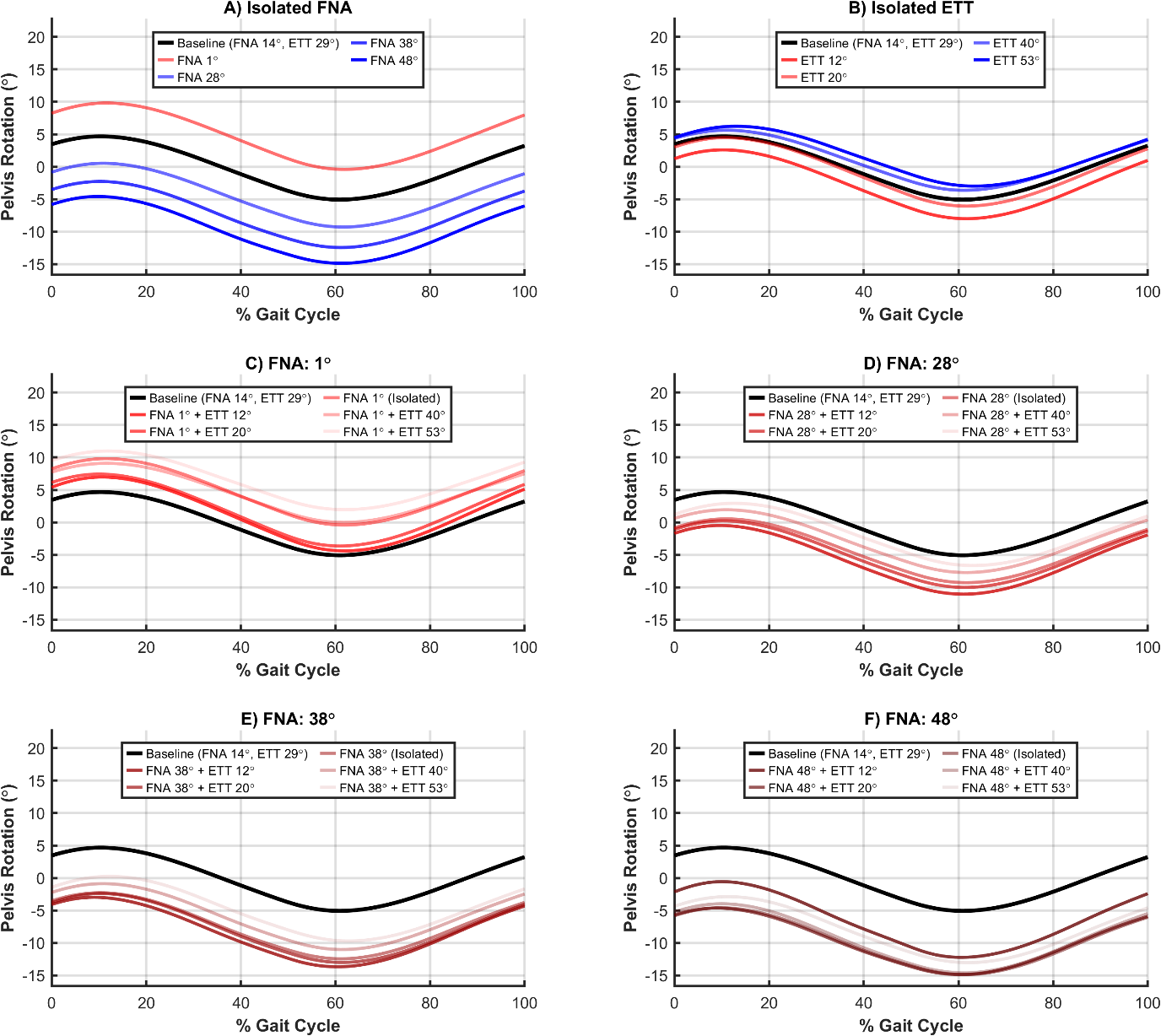

**Supplementary Figure 8** Pelvis internal-external rotation (positive values indicate internal rotation, counterclockwise rotation, of the pelvis relative to the global reference frame) over the gait cycle for isolated and combined femoral neck anteversion (FNA) and external tibial torsion (ETT) conditions. The first row shows isolated FNA (A) and ETT (B) conditions. The bottom two rows (C-F) show fixed FNA angles with varied ETT angles.

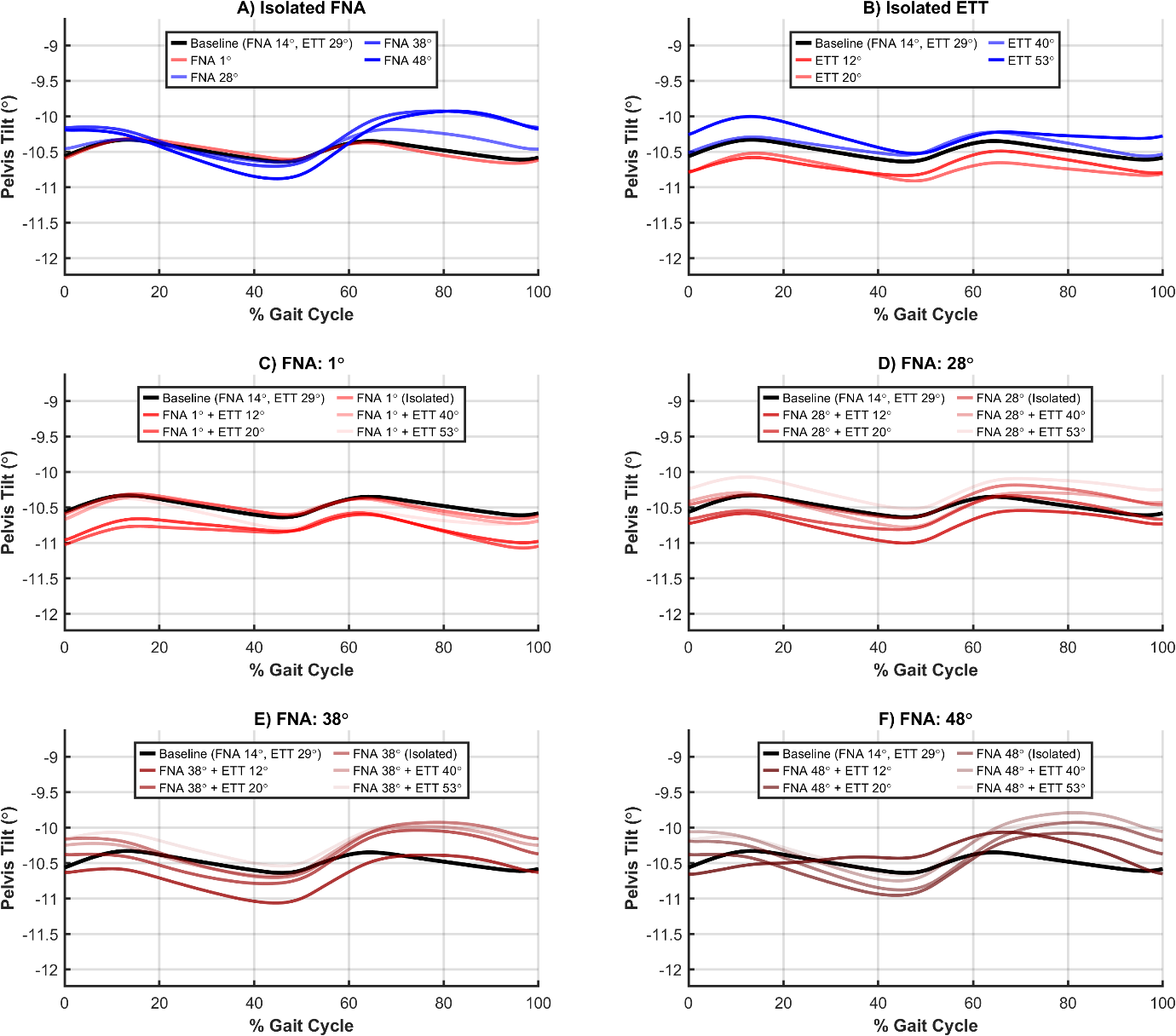

***Supplementary Figure 9*** Pelvis tilt (negative values indicate forward tilt, clockwise rotation, of the pelvis relative to the global reference frame) over the gait cycle for isolated and combined femoral neck anteversion (FNA) and external tibial torsion (ETT) conditions. The first row shows isolated FNA (A) and ETT (B) conditions. The bottom two rows (C-F) show fixed FNA angles with varied ETT angles.

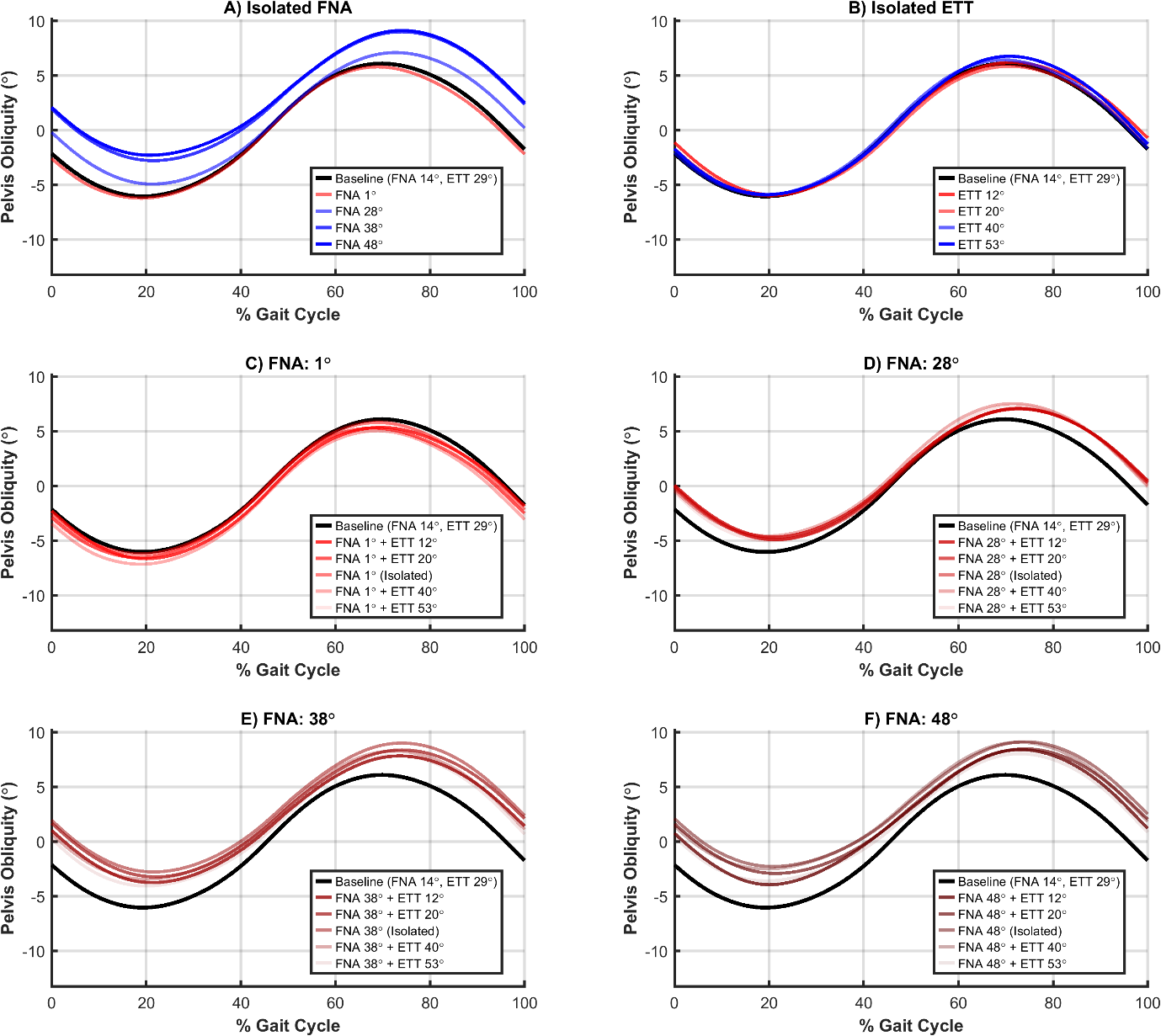

***Supplementary Figure 10*** Pelvis obliquity (positive values indicate contralateral pelvic drop, anticlockwise rotation, of the pelvis relative to the global reference frame) over the gait cycle for isolated and combined femoral neck anteversion (FNA) and external tibial torsion (ETT) conditions. The first row shows isolated FNA (A) and ETT (B) conditions. The bottom two rows (C-F) show fixed FNA angles with varied ETT angles.

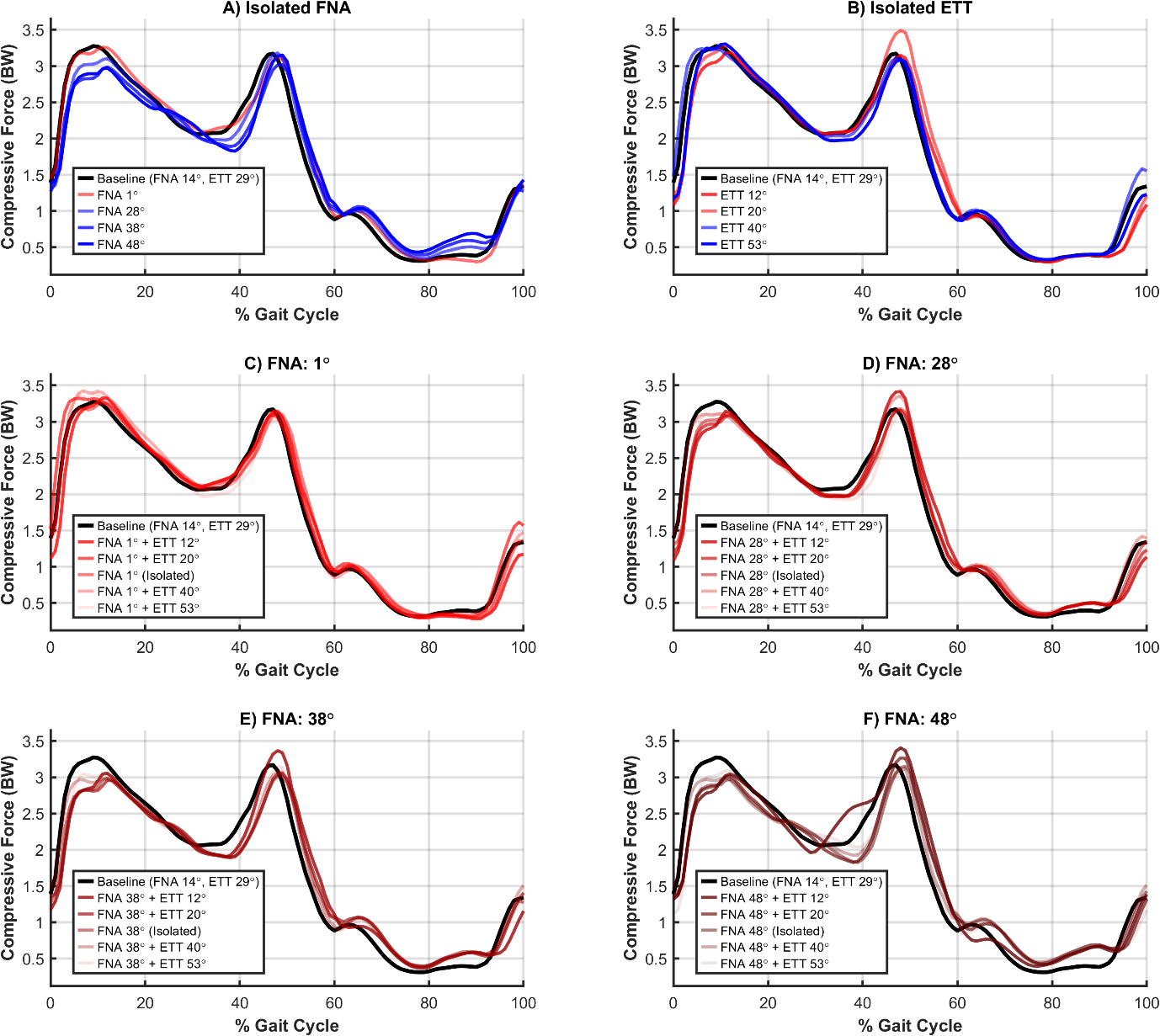

**Supplementary Figure 11** Compressive hip joint contact force normalised to body weight (BW) over the gait cycle for isolated and combined femoral neck anteversion (FNA) and external tibial torsion (ETT) conditions. The first row shows isolated FNA (A) and ETT (B) conditions. The bottom two rows (C-F) show fixed FNA angles with varied ETT angles.

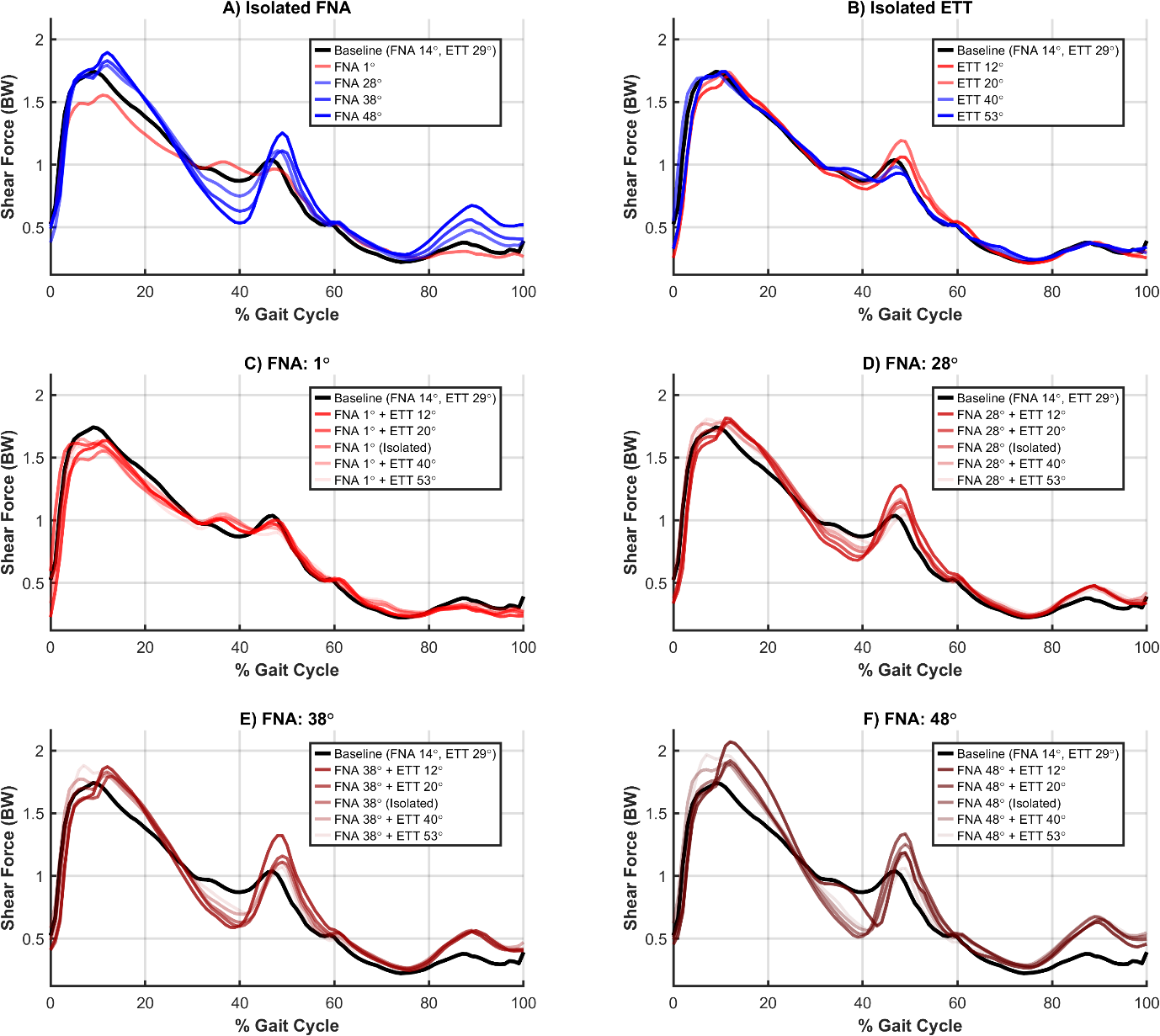

**Supplementary Figure 12** Shear hip joint contact force normalised to body weight (BW) over the gait cycle for isolated and combined femoral neck anteversion (FNA) and external tibial torsion (ETT) conditions. The first row shows isolated FNA (A) and ETT (B) conditions. The bottom two rows (C-F) show fixed FNA angles with varied ETT angles.

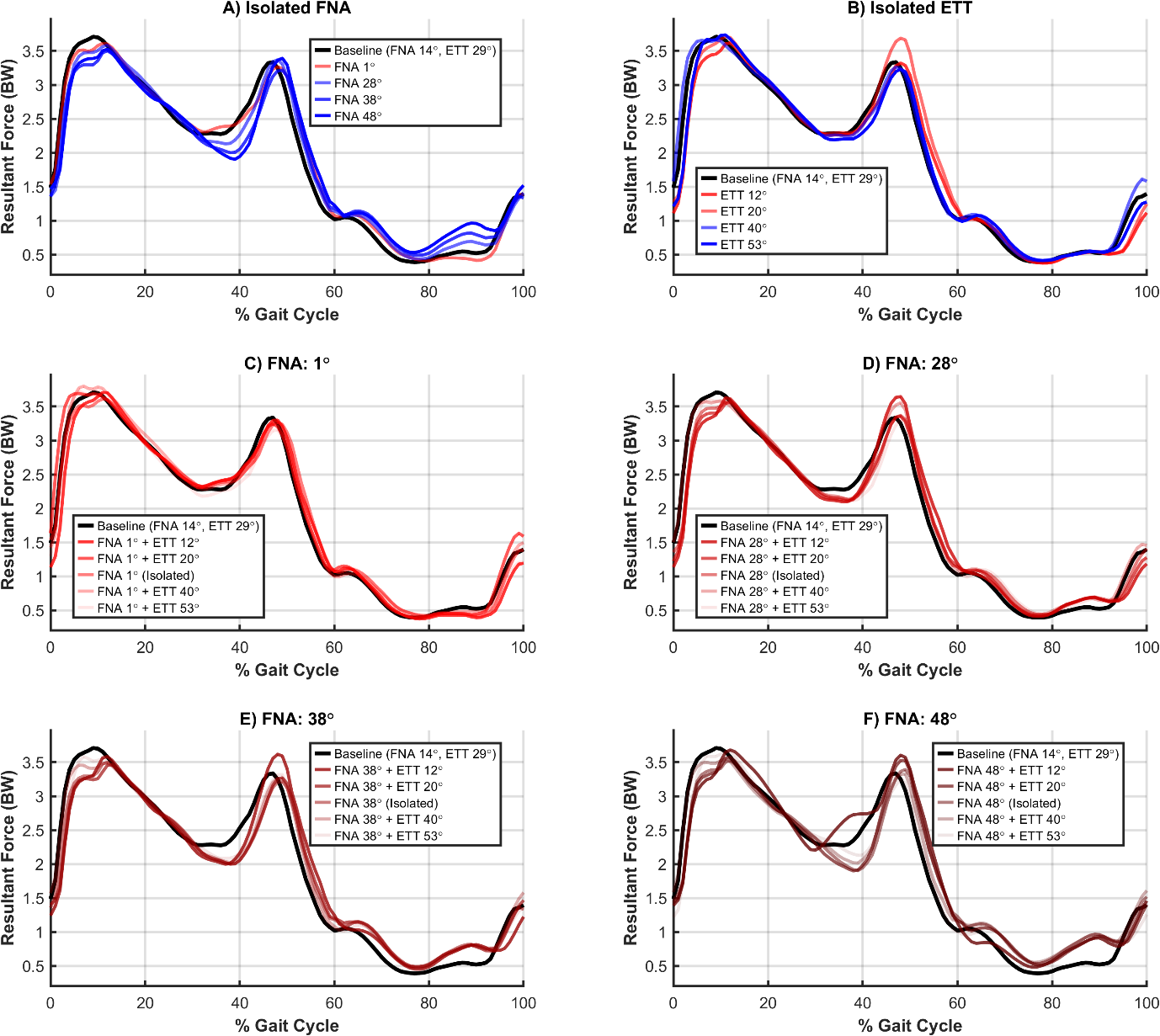

**Supplementary Figure 13** Resultant hip joint contact force normalised to body weight (BW) over the gait cycle for isolated and combined femoral neck anteversion (FNA) and external tibial torsion (ETT) conditions. The first row shows isolated FNA (A) and ETT (B) conditions. The bottom two rows (C-F) show fixed FNA angles with varied ETT angles.

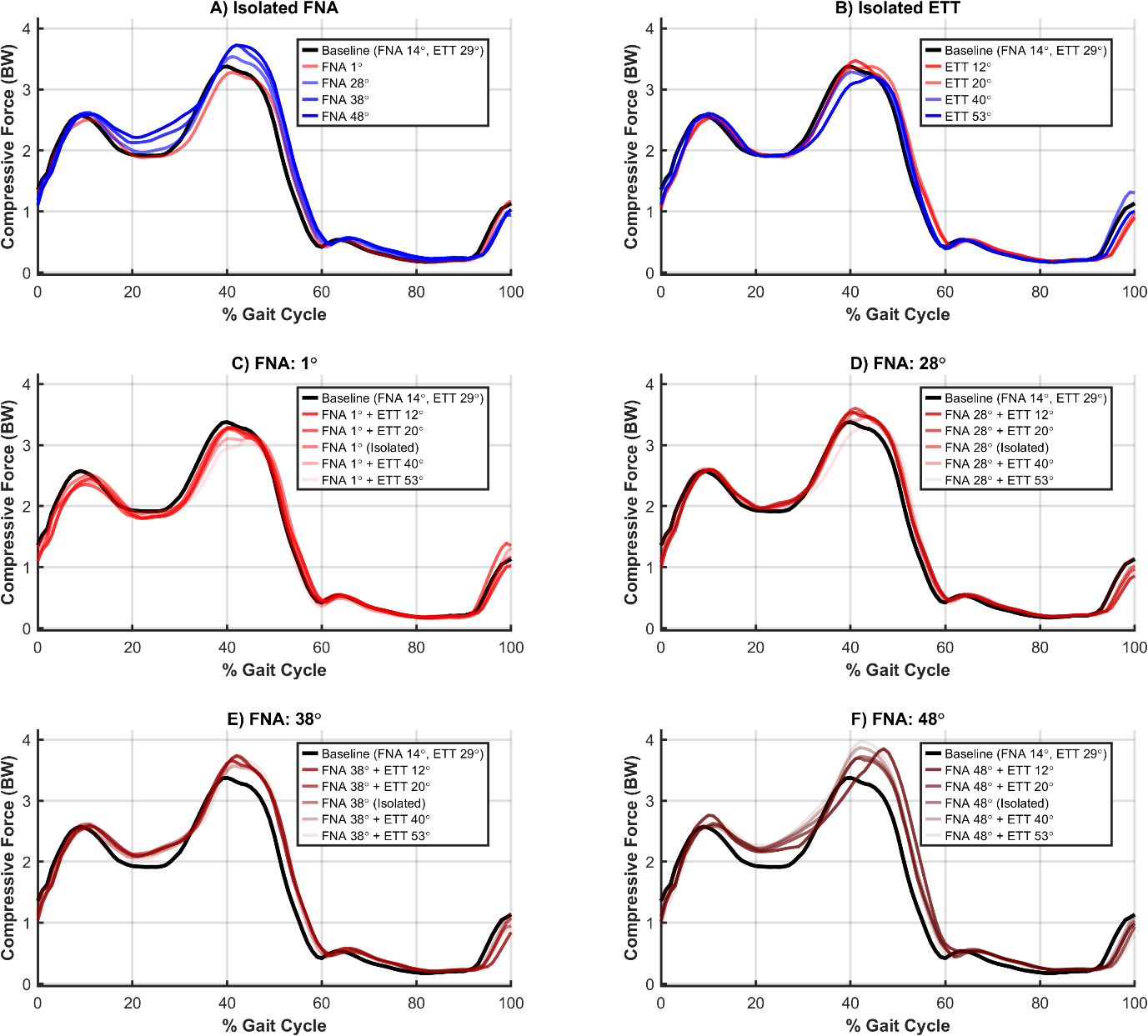

**Supplementary Figure 14** Compressive knee joint contact force normalised to body weight (BW) over the gait cycle for isolated and combined femoral neck anteversion (FNA) and external tibial torsion (ETT) conditions. The first row shows isolated FNA (A) and ETT (B) conditions. The bottom two rows (C-F) show fixed FNA angles with varied ETT angles.

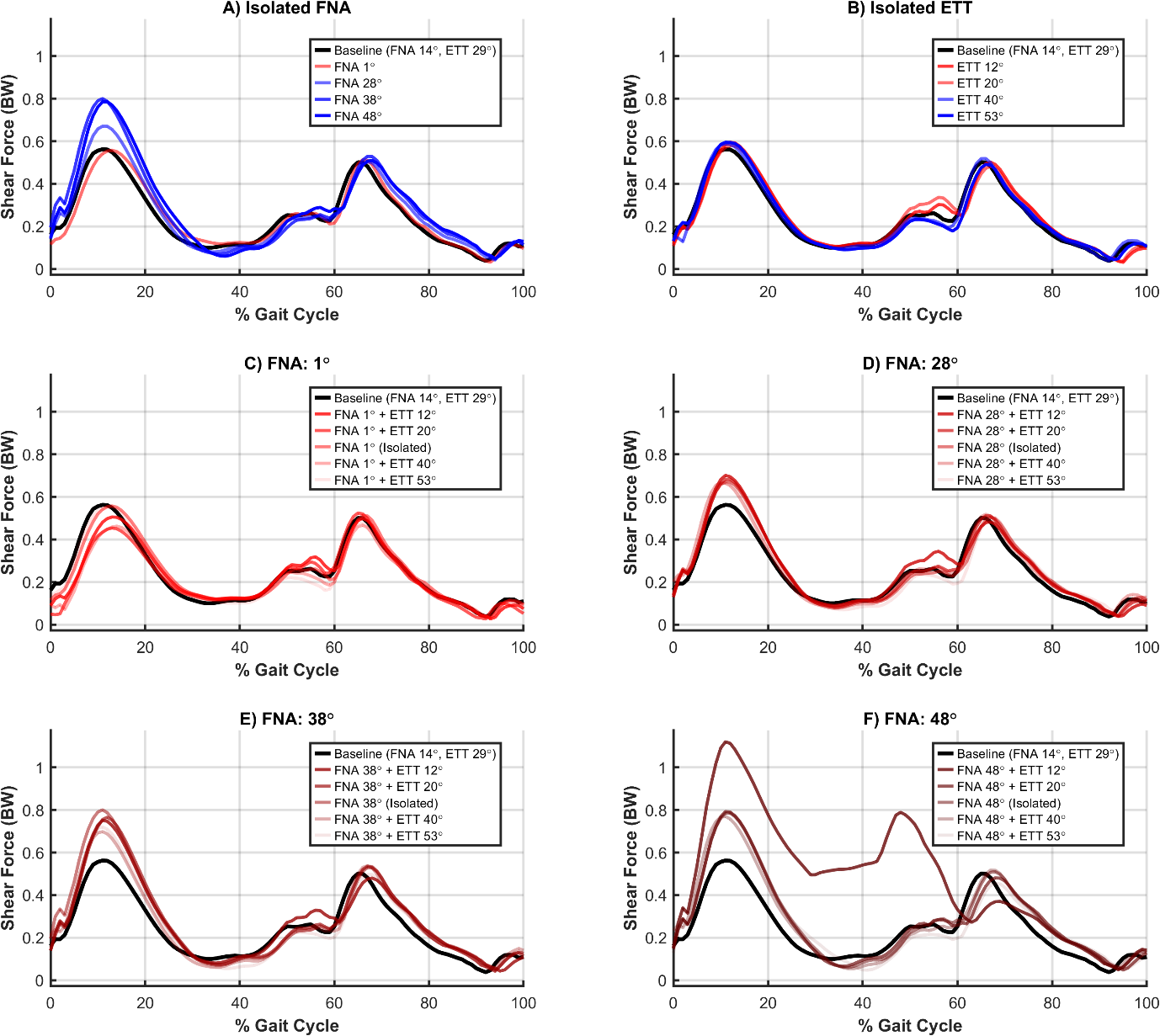

**Supplementary Figure 15** Shear knee joint contact force normalised to body weight (BW) over the gait cycle for isolated and combined femoral neck anteversion (FNA) and external tibial torsion (ETT) conditions. The first row shows isolated FNA (A) and ETT (B) conditions. The bottom two rows (C-F) show fixed FNA angles with varied ETT angles.

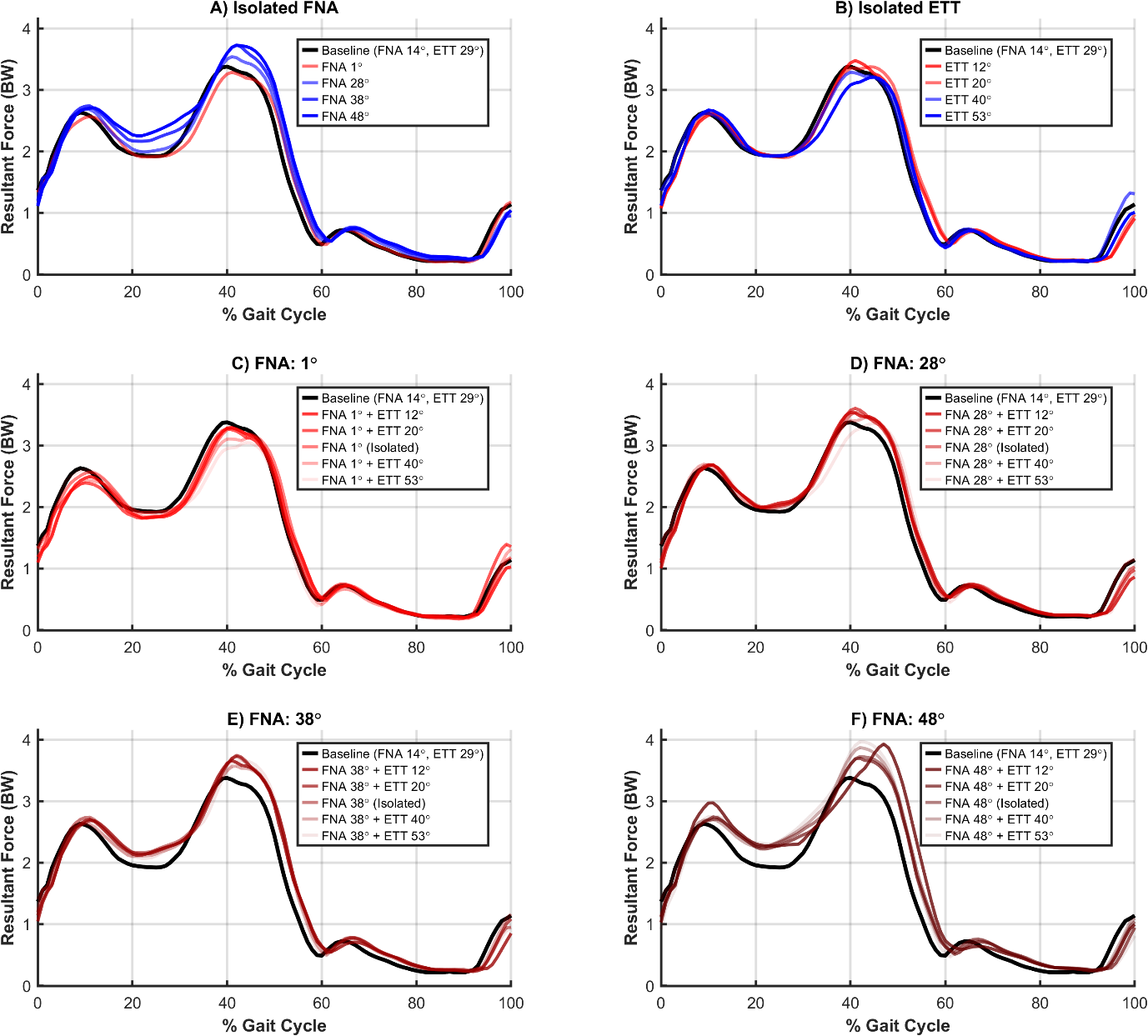

**Supplementary Figure 16** Resultant knee joint contact force normalised to body weight (BW) over the gait cycle for isolated and combined femoral neck anteversion (FNA) and external tibial torsion (ETT) conditions. The first row shows isolated FNA (A) and ETT (B) conditions. The bottom two rows (C-F) show fixed FNA angles with varied ETT angles.
